# DISPERSAL BEHAVIOR RATHER THAN DISPERSAL MORPHOLOGY CREATES SOCIAL POLYMORPHISM IN FORMICA ANTS

**DOI:** 10.1101/2025.06.27.661689

**Authors:** Sanja Maria Hakala, Ilya Belevich, Eija Jokitalo, Perttu Seppä, Heikki Helanterä

**Affiliations:** Centre of Excellence in Biological Interactions, Organismal and Evolutionary Biology Research Programme, Faculty of Biological and Environmental Sciences, University of Helsinki, Finland; Tvärminne Zoological Station, University of Helsinki, Finland; Department of Ecology and Evolution, Faculty of Biology and Medicine, University of Lausanne, Switzerland; Electron Microscopy Unit, University of Helsinki, Finland; Ecology and genetics research unit, University of Oulu, Finland

**Keywords:** Sex biased dispersal, resource allocation, social evolution, Hymenoptera, Formicidae, polydomy, polygyny, supercoloniality

## Abstract

1. Dispersal evolution and social evolution are interlinked. Dispersal is necessary for avoiding kin competition and inbreeding, but limited dispersal also allows beneficial social interactions with kin. In ants, a correlation between poor dispersal and complex societies, where a big proportion of queens are philopatric, is well documented, but the underlying causal mechanisms are not clear.
2. In this study we investigate the dispersal ability of six *Formica* species that vary in their colony queen number and nest founding mode, from three different subgenera. Investigating resource allocation in the bodies of young queens and males allows us to analyze the evolutionary causalities between dispersal ability and social organization. We measured the body ratios including wing-muscle ratio; glycogen, triglyceride and protein resources with colorimetric assays; and microscopic wing muscle structures with transmission electron microscopy, with an overall sample size of 1515 individual males and queens.
3. Our results suggest that the physical condition of individuals does not strongly correlate with the dispersal patterns of the species, contrary to assumptions based on earlier studies in both ants and other insects. There was still a minor effect of social organization on male wing-muscle ratio, which is an interesting case of sexual coevolution of dispersal traits: it is likely that the queen philopatric behavior is reflected in male behavior and consequently in male morphology – even when our overall results support the hypothesis that ant dispersal is male biased regardless of social organization. Further, our intraspecific analyses in two of the six species reveal different patterns in their flight abilities in connection to their social organization, further pointing towards a mismatch between dispersal behavior and ability.
4. In *Formica* queens, philopatry seems to be a behavioral trait of the individuals rather than a consequence of colony-level resource allocation into dispersal ability, pointing towards a “behavior first” evolutionary route. The queens may selfishly choose not to disperse even when their society provides them with the resources for it. This raises new questions about conflicts over dispersal in these highly social systems.

## INTRODUCTION

In natural populations, local resource competition and potential habitat destruction require that organisms move to new locations in order to survive on evolutionary time scales (Van Valen 1971). Resource competition among relatives is especially harmful, as it decreases inclusive fitness, which reinforces selection for dispersal – even in social organisms that simultaneously also benefit from interactions with their relatives (Hamilton and May 1977, West et al. 2002). Dispersal also allows individuals to avoid breeding with their kin, which often selects for sex-biased dispersal (Bengtsson 1978). Evolving and maintaining efficient dispersal strategies is crucial for all organisms.

Social insect colonies can be seen as superorganisms that are bound to their sedentary nesting sites and often only disperse through their mobile sexual castes, the young queens and males produced in the colonies (Helanterä 2016, Hakala et al. 2019). For most social insects, dispersal by wing is an inseparable element of their reproductive strategy (Boomsma et al. 2005), and dispersal shapes the evolution of the sexual castes. In most ant species, the young queens and males leave their natal colonies for mating flights, after which the males die and the queens found new colonies (Bourke and Franks 1995). Although the queens spend most of their adult lives within their colonies, the strong selection pressures during dispersal are a major driver of their evolution (Helms 2018). Flight-related trade-offs and resource allocation affect the evolution of the queens (Helms and Kaspari 2015), and thus the evolution of the colonies they found. The adult males, who are not directly involved in the colony social life, have an indirect effect through mating and dispersal strategies that coevolve with the queen strategies (Heinze and Tsuji 1995, Peeters and Aron 2017, Hakala et al. 2019)

In some ant species, all daughter queens disperse and found new independent colonies, whereas in others a proportion of them stays in their natal colonies as extra queens resulting in polygynous colonies. In such colonies, queens reproduce cooperatively and share resources, including the worker force (Boomsma et al. 2014). In some ant taxa, polygyny is associated with polydomous colony organization, where the society (colony) occupies several interconnected nests. Such a strategy allows more effective resource acquisition and often also a yet higher number of queens and thus an even larger colony size (Debout et al. 2007). In some taxa, polygynous and polydomous societies may develop into extremely large network colonies with thousands – or even more – of egg-laying queens, when they are referred to as supercolonies (Helanterä et al. 2009, Helanterä 2022). This kind of continuum from simple (monogynous and/or monodomous) to more complex societies (polygynous and polydomous) even among closely related species and among different colonies of the same species (Hölldobler and Wilson 1977), offers an informative study system for inferring the coevolution of social organization and dispersal.

Correlation between limited dispersal and polygynous and polydomous societies in ants is well documented with gene flow analyses in several taxa (Seppä 2008). Ant community analyses also support this: species with polydomous social organization seem to be poorer at reaching isolated areas (e.g. Mabelis 1994; Sorvari 2017). However, the exact evolutionary reasons and causal mechanisms for this correlation have not been assessed in detail. The *Formica* ants are known to carry a supergene associated with the colony queen number (Brelsford et al. 2020), but curiously the most polygynous and polydomous species may have lost the allele associated with polygyny in other species (Lagunas-Roblesa et al. 2024, Sigeman et al. 2024). Regardless of the genetic architecture, the functional mechanisms creating the largest polydomous nest networks are still unclear. It is obvious that dispersal behavior differs dramatically between monodomous and highly polydomous social organization – philopatric queens are an inbuilt characteristic that create the polydomous and supercolonial ant societies. One of the main unanswered questions is, whether the observed difference in queen dispersal is due to difference in the dispersal ability and morphology or whether it is an active behavioral choice.

Ant males have been studied much less than the two female castes, and data on ant male dispersal is sparse (Boomsma et al. 2005, Shik et al. 2013, Hakala et al. 2019). As ant dispersal coevolves with their mating strategies, strategies of both sexes must be studied. It seems likely that ant dispersal is generally male biased due to higher evolutionary restrictions and trade-offs affecting the queens, but there is also a bias towards studying species with more complex social organizations (Hakala et al. 2019). In *Formica* ants, for instance, dispersal seems to be especially strongly male biased in polygynous and polydomous species (Sundström et al. 2005), but this pattern does not necessarily hold in a strictly monodomous species (Johansson et al. 2018). Studying a wide range of species with varying social organizations is required to assess whether male bias is a general characteristic of the superorganism dispersal strategies.

In this study, we investigate the dispersal ability of three species pairs of *Formica* ants, from three different subgenera. The contrast of monodomous social organization with high levels of polydomy in each of the three species pairs provides an excellent model for studying the evolution of dispersal ability and social organization. The queens of all these species have wings, but it is unclear if the ability to fly and disperse is weakened in the predominantly polydomous species, where the queens regularly do not fly. Our main hypothesis is that philopatric queen behavior is reflected in their morphology, so that the polydomous species that disperse less have smaller allocation to dispersal traits. Additionally, we expect male dispersal traits to be stronger overall than the queen traits. However, the hypotheses for male dispersal traits in connection to social organization are difficult to formulate due to lack of studies on ant male dispersal. It is unclear, whether and how the change in social organization affects mating competition or the costs of inbreeding for the haploid males and diploid queens, and how this in turn affects the coevolving dispersal strategies of the two sexes. We present two opposing hypotheses: 1. The selection pressures for both sexes are similar, and thus also male dispersal traits are weaker in polydomous than in monodomous species. 2. When female dispersal strategy shifts towards philopatry in polydomous societies, selection for male dispersal gets stronger, which leads to polydomous males evolving stronger dispersal traits.

## MATERIALS AND METHODS

### SPECIES AND POPULATIONS

We sampled six *Formica* species from three different subgenera: *F. pratensis* and *F. aquilonia* from *F. rufa sensu stricto*, *F. exsecta* and *F. pressilabris* from *Coptoformica*, and *F. fusca* and *F. cinerea* from *Serviformica*, forming three species pairs, each with a species with predominantly simple monodomous societies and a species with complex, highly polygynous and polydomous societies (Table 1). In *Formica* ants, polydomous colonies can generally grow into supercolonies consisting of even thousands of nests (Markó et al. 2012). The two *Coptoformica* species have both types of societies in our study area, and we sampled both for intraspecific analyses. Also *F. cinerea* has been reported to have monodomous colonies in the area (Goropashnaya et al. 2001), but we were not able to find enough of them to include this species for intraspecific analyses. For the main analysis including all the species, only the typical monodomous populations of *F. exsecta* were included, leaving out the supercolony atypical for the area. Similarly, only the well-established large polydomous population of *F. pressilabris* is included in the among species analysis, leaving out the monodomous population inhabiting a clear-cut forest area and consisting of small nests only. All the study populations are located in the vicinity of Tvärminne Zoological Station in southern Finland.

**Table 1:**
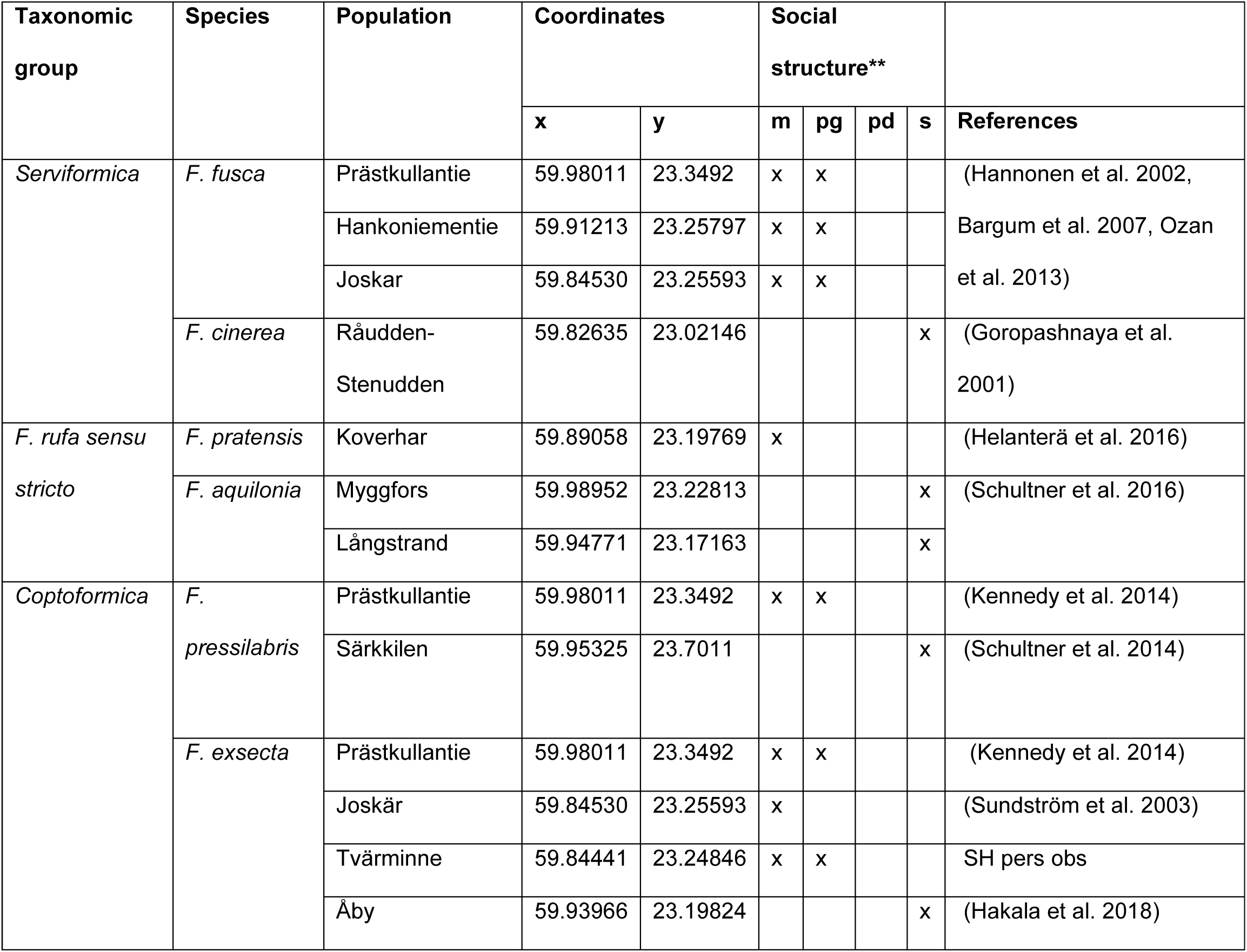
The sampled *Formica* species and the social organization of the populations studied. ** The social structure: m=monogyny (one queen in a colony), pg=polygyny (several queens in a colony), pd=polydomy (several nests in a colony), s=supercoloniality (huge polygynous and polydomous colonies). The social structures of these populations are analysed in the referenced previous studies.

### FIELD SAMPLING

We monitored natural field colonies of the six *Formica* species for maturation of sexual brood by gently excavating the mounds. The *F. rufa* group species were sampled solely from the colonies in the field to ensure uninterrupted development. We sampled young males and virgin daughter queens when they emerged on top of the nest. No individuals were sampled from within the nest, as their age and maturity could not be assessed without this behavioral signal. *Serviformica* species have underground colonies that cannot be monitored for exact timing of emergence of the sexuals without serious disruption of the nest structures. Thus, for these species, nest material, workers and brood were sampled from the field colonies when the brood was in late pupal stage and brought to the laboratory for maturation. The sexual individuals were subsequently sampled from the laboratory nests when they emerged on top of the nest, as described above. *Coptoformica* individuals were mostly sampled from the field colonies, but the sample set was supplemented with laboratory-grown individuals to ensure high enough sample size. *Formica pressilabris* queens were never observed on top of their nests in the field colonies, and therefore they are all laboratory-grown. Sampling was mostly done in 2016, but the data for *F. fusca* and *F. cinerea* was supplemented in 2017 to ensure high enough sample sizes. The spermathecae of c. 20 daughter queens from several different colonies of each of the study species were dissected to ensure that they were unmated.

Nests were housed in plastic boxes coated with fluon to prevent ants climbing out, and covered with a cloth to prevent males and queens flying out. The nests were watered daily and maintained with Bhatkar & Whitcomb diet (1970) in room temperature and natural light cycle. The laboratory rearing took place in Tvärminne Zoological Station (for 2016 samples) or in the Viikki Plant Growth Facilities in Helsinki (for 2017 samples).

After sampling, the individuals were kept on ice to minimize water loss and usage of metabolic resources during the transfer. Each individual’s (n=1515) wet mass was measured in three parts separately: head, thorax (with legs but without the wings), and abdomen. After weighing, specimens were snap frozen in liquid nitrogen and stored in -80 °C for muscle size and metabolic analyzes. As the size of these species varies considerably, thorax mass and abdomen mass ratios were calculated and used for further analyses instead of the absolute mass.

### BODY PROPORTIONS AND FLIGHT MUSCLE RATIO

The dispersal ability of insects is a sum of several physiological and morphological traits, and a subject to clear trade-offs. As dispersal happens by wing, size of the direct and indirect flight muscles is one of the most important factors (Dudley 1999). In several insects, the thorax mass / body mass ratio (hereafter thorax ratio) has been identified as a key predictor for flight ability and dispersal (e.g. Srygley and Chai 1990; Thomas et al. 1998; Berwaerts et al. 2002; Steyn et al. 2016), also including ants (Helms and Kaspari 2014, Helms and Godfrey 2016). However, in addition to just flight muscles, ant females have strong neck muscles inside their thoraxes, taking different amounts of space in different species (Keller et al. 2014). Additionally, ant queens can histolyse their flight muscles (Janet 1907, Jones 1979, Hölldobler and Wilson 1990), which means that their presence in the thorax cannot be guaranteed without direct observation. Thus, the thorax ratio cannot be assumed to represent the flight ability in a straightforward way across different ant species. Therefore we opted for measuring the muscle size directly by using the wing muscle mass/ body mass ratio, which correlates well with flight performance across a wide range of animal phyla (Marden 1987; 1989).

Flight muscle ratios (hereafter muscle ratio) were measured for a total of 627 specimens by dissection as described by Marden (1987). Individual specimens were weighed alone, except for the smallest *F. pressilabris* and *F. exsecta* males, where males were pooled within the nests to improve the repeatability of weighing. For these specimens, the final muscle sizes are nest averages.

### TRANSMISSION ELECTRON MICROSCOPY

In addition to muscle size, also the structure of the muscles affects flight performance (Marden 2000). The myofibril and mitochondria densities correlate with flight performance (Sohal et al. 1972, Fernandes et al. 1991, Rauhamäki et al. 2014). The mitochondrial inner membranes host the cytochrome c oxidase enzyme that catalyzes the reduction of oxygen to water in cellular respiration, and affects directly the metabolic rate and thus the flight performance of the individuals (Niitepõld et al. 2009, Rauhamäki et al. 2014). In ants, detailed studies on the microscopic muscle structures are rare, but a study in the socially polymorphic *Formica truncorum* showed that queens from polydomous colonies have slightly fewer mitochondria in their muscles than queens from monodomous colonies - suggesting poorer flight ability (Johnson et al. 2005). Another aspect of muscle structures that is especially important for ants is the possibility of flight muscle histolysis. It is usually thought to happen only in queens after the dispersal flight (Hölldobler and Wilson 1990), but this has not been widely tested. In many other insects, flight muscles can be histolysed also before flight, and damage from histolysis may render the muscles useless even if they are still present (Fairbairn and Desranleau 1987).

To measure these muscle structure variables, nine additional individuals from three different nests in each species and both sexes (n=108) were prepared for transmission electron microscopy analysis of the flight muscles. After weighing the individuals, their thoraxes were dissected to allow the fixation buffer to penetrate the muscle tissues. Legs and leg and neck muscle tissues were removed, and the petiole was cut off. The thorax was ventrally opened, but flight muscle tissue was kept intact inside the thorax. The specimen was submerged in fixation buffer containing 2% glutaraldehyde, 100mM Na-cacodylate buffer pH 7.4, 150mM saccharose, first for 2h at room temperature and subsequently for 22h at +4C. After fixation, samples were stored 1-2 months in a solution containing 2% formaldehyde, 100mM Na-cacodylate buffer pH 7.4 and 150mM saccharose.

Further sample handling was done in the Electron Microscopy Unit in the Institute of Biotechnology, University of Helsinki. The samples were post-fixed using 2% non-reduced OsO_4_ (EMS, Hatfield, PA) in H_2_O for 2h at room temperature; dehydrated in ethanol gradient (50%, 70%, 96%, 100%, 100%, 15 min. each) and 99.5% acetone (Sigma) for 30 min; infiltrated into low-viscosity resin (TAAB, LV Medium) using the following resin: acetone ratios: 1:1 (2h), 2:1 (2h), 1:0 (o/n), 1:0 (4h). The resin blocks were cured in 60°C overnight. Mounting orientation of the final sample blocks was adjusted for observing cross sections of the myofibrils, verified by roundness of cross-sections of myofilaments.

For morphometric analyses for each specimen, systematic random sampling was used to obtain 10-14 images from different cells at the magnification of 2500x (single image area = 69.2 µm^2^, n=1 240) with Jeol JEM-1400 transmission electron microscope, equipped with Gatan Orius SC 1000B bottom mounted CCD-camera. The collected 16-bit TEM images were stored in DM4 format, and processed using the software Microscopy Image Browser (MIB) (Belevich et al. 2016). The contrast was normalized such that the mean intensity for each image was set to 24000 and standard deviation of the intensity histogram to 3000 counts. After that the images were converted to the 8-bit format using intensity 12000 as a black point and 42000 as white point. To assess differences in mitochondrial and myofibril size and density, their profiles were manually traced using MIB and an interactive pen display. The mitochondria profiles were analyzed into two categories: 1) complete profiles and 2) the profiles clipped at the edges of the images. To assess the total mitochondria and myofibril areas (µm^2^) per image, we used Stereology tool of MIB with the sampling grid size of 50 x 50 nm.

The analysis of individual cell organelles was done using a custom-made plugin of MIB. For each image the following properties were extracted for each complete myofibril profile (n=9 757): *CentroidX, CentroidY, TotalArea, minDiameterAverage* (calculated by using ultimate erosion to find the middle line of the organelle profile; measuring the distance to the closest edge for each point of the middle line; converting the collected values to the diameter by multiplying them by 2 plus pixel size; and finally averaging the measurements); and from each complete mitochondrion profile (n=27 633): *CentroidX, CentroidY, TotalArea, ThresholdedArea, RatioOfAreas, minDiameterAverage*. For estimation of degradation of mitochondria we used an empirical threshold value of 140 and calculated the ratio of thresholded vs. original area for each mitochondrion.

To analyze the organization of myofibrils in our images, distances among the centroids of the closest complete myofibrils in our images were calculated by Delaunay triangulation (‘delaunay’ function in Matlab), excluding the distances on the outer border (freeBoundary option). The bordering distances do not represent true distances to the closest organelles as the neighboring organelles were clipped with the image edges. To get a measure independent from the size of the myofibrils, we analyze the relative standard deviation (=coefficient of variation) of their distances for each image. The smaller the CV value, the more evenly organized the myofibrils are.

To assess the shape of mitochondria in our images, a value for mitochondrion shape was calculated as follows:

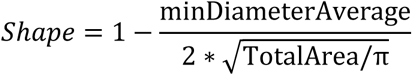

Thus, a completely round mitochondrion profile shape gets a value of 0, and the closer the value is to 1, the longer the profile shape is. Long mitochondria profile shapes (*Shape* > 0.5) indicate the existence of elongated, tubular mitochondria, although their amount in the cross-section images will be low due to the way elongated mitochondria align with the myofibrils (Yu et al. 2003).

### COLORIMETRIC ASSAYS FOR GLYCOGEN, TRIGLYCERIDE AND PROTEIN ANALYSIS

In ants, the main fuel for flight is glucose stored in the flight muscles and fat body as glycogen (Peakin 1964, Toom et al. 1976, Passera et al. 1990, Wegener 1996). In many insects, also fatty acids are used as a biochemical fuel for prolonged flight (Chino and Downer 1982, Wegener 1996), but in ants they seem to be only used by queens as an energy source during the nest founding period (Keller and Passera 1989, Wheeler and Buck 1996). Males have been reported to have negligible amounts of stored fat (Boomsma and Isaaks 1985). Along with fatty acids, queens also store proteins to be used as energy source during nest founding, and these storage proteins form a substantial proportion of their total protein amount (Wheeler and Buck 1995, 1996, Wheeler and Martínez 1995). Several previous studies show that these biochemical traits correlate with flight and nest founding strategies in ants, both within-species and among-species: Queens participating in the mating flights and founding their nests independently have higher amounts of glycogen, fat and storage proteins, compared to queens not participating in the mating flights but instead founding their nests dependently with workers from their natal nest (Keller and Passera 1989, Passera and Keller 1990, Sundström 1995, Hahn et al. 2004). Similar results have been shown for the male glycogen amounts, although there are substantially less studies done on them (Passera and Keller 1990, Sundström 1995).

Glycogen, triglyceride and protein concentrations were measured for 700 different individuals from the same nests that were used for muscle size measurements. All protocols were adjusted from Tennessen et al. (2014) to allow for measuring all concentrations from the same individuals, in 384 well plate format to allow small sample volumes. Individual ants were measured for most species, with the exception of *F. pressilabris* males and females, and small *F. exsecta* males that were analyzed pooled with the other individuals collected from the same nest to ensure reliable results. Body weight of 10mg was used as a cut-off for pooling. The sample used in these measurements contained the thorax and abdomen. The head was not used as eye pigments may interfere with the measurement, and the head does not contain storage resources.

The deep-frozen samples were homogenized with Qiagen Tissuelyser II machine and metal beads prior to adding buffer. To allow comparing the results of different sized individuals, the samples were standardized by their wet weight by using 10ul BPST buffer/ 1 mg of mass of the whole individual. Glycocen concentrations were analyzed by breaking the glycogen to glucose with amyloglucosidase and comparing the obtained total glucose concentration to the free glucose concentration of each sample as a paired assay with Sigma Aldrich Glucose (GO) Assay Kit. The triglyceride concentrations were analyzed with Sigma Aldrich Serum Triglyceride Determination Kit. Protein concentrations were analyzed with Sigma Aldrich Bradford Reagent. The absorbances were measured with the Enspire multimode plate reader.

### STATISTICAL ANALYSES

The data were analyzed in R with the package lme4 (Bates et al. 2014), and estimates for p-values were obtained with the package lmerTest (Kuznetsova et al. 2017). Linear mixed models (LMM) were used for the body proportion and biochemical data. In all these analyses, the measured variable was used as a response variable, and the social organization (monodomous/ polydomous), sex, and their interaction were used as fixed effects, except for the triglyceride measurements where there are data only for the queens and thus only the social organization is used as fixed effect. The species and nest were used as nested random effects. The similarity of variances between males and females, as well as between the two social types, were tested with an F-test. As the variance of these data differ between the inspected classes in some cases, the reliability of LMM results was checked with robust LMM models that give more weight to samples with values closer to the mean, with the package robustlmm (Koller 2016).

Similar LMM analyses on body proportions and biochemical measurements were repeated for *F. exsecta* and *F. pressilabris* with additional samples from the divergent social organization (polydomous for *F. exsecta*, and monodomous for *F. pressilabris*) to analyze within-species variation. In these analyses the nest was used as a random effect. The polydomous *F. exsecta* population produces an extremely male-biased sex ratio, and thus our sample size for polydomous *F. exsecta* queens is very small (N=7 from a single nest). All of them were used for the muscle size analysis, and thus the effect of social organization on the biochemical resources was not analyzed for *F. exsecta* queens.

The microscopical muscle structure data were analyzed similarly with either LMMs or with generalized linear mixed models (GLMM) with gamma distribution and an identity link function. As most mitochondria profiles get small values of degradation with our initial thresholding, the initial values of degradation are not especially informative. Thus, the presence of high values of degradation (>10% of mitochondria profile area) was analyzed with a GLMM with binomial distribution and a logit link function. In all data for individual cell organelles the nested random effect structure was as follows: species/nest/individual/image. For the image level measurements (total mitochondria/myofibril area and the CV of distances among myofibrils) the random effect structure was species/nest/individual.

All models were validated as instructed by (Zuur 2016). Benjamini-Hochberg procedure (1995) was used to correct p-values of all of the LMMs and GLMMs to decrease the false discovery rate arising from high number of measurements and corresponding statistical tests. False discovery rate of 0.05 was used in the correction.

The microscopic muscle structure data were combined into a single data set by calculating image means for the measured variables. Correlations among the measured variables for both sexes separately were visualized with the R package corrplot (Wei and Simko 2017). The associations between different measurements were inspected with Pearson’s product moment correlation coefficient with the significance level p = 0.01. The dataset was centered, scaled and transformed (Box and Cox 1964), and linear discriminant analyses (LDA) were performed with the R package MASS (Venables and Ripley 2002) for males and queens separately to investigate whether the six species or the two social types differ overall by their dispersal traits. As none off the measured microscopical traits were too strongly intercorrelated (cut-off correlation coefficient 0.7) they were all kept in the analyses (Appendix 3: Figure 5). The LDA accuracy was tested by ten-fold cross-validation with the R packages *caret* (Kuhn 2008).

The body ratio and biochemical data were similarly combined into a single data set by calculating nest means for the measured variables for all the main social type nests of all the six species. Correlations were analyzed and LDA performed as above. For males the LDA included the total body weight, abdomen ratio, muscle ratio, protein concentration and glycogen concentration. Thorax ratio was not included due to a strong correlation with abdomen ratio (correlation coefficient -0.85, Appendix 2: Figure 2). For the queens the LDAs included abdomen ratio, muscle ratio, protein concentration, glycogen concentration and triglyceride concentration. Total body weight and thorax ratio were not included due to a strong correlation with abdomen ratio (correlation coefficients 0.70 and -0.87, respectively, Appendix 2: Figure 2).

## RESULTS

All results for all the measured variables for each species, social organization and sex are summarized in tables in Appendix 1.

### BODY PROPORTIONS AND FLIGHT MUSCLE RATIO

The muscle ratios were higher in males (species averages between 0.23 - 0.31) than in queens (species averages 0.15 - 0.25) (Figure 1, Table 2). There was a trend of male muscle ratios being smaller in polydomous species than monodomous species, with no effect in queens, seen in the significant interaction in our models (Figure 1, Table 2). Polydomous species also had larger variance of the muscle ratio than the monodomous species (Appendix 2: Table 1).

**Figure 1:**
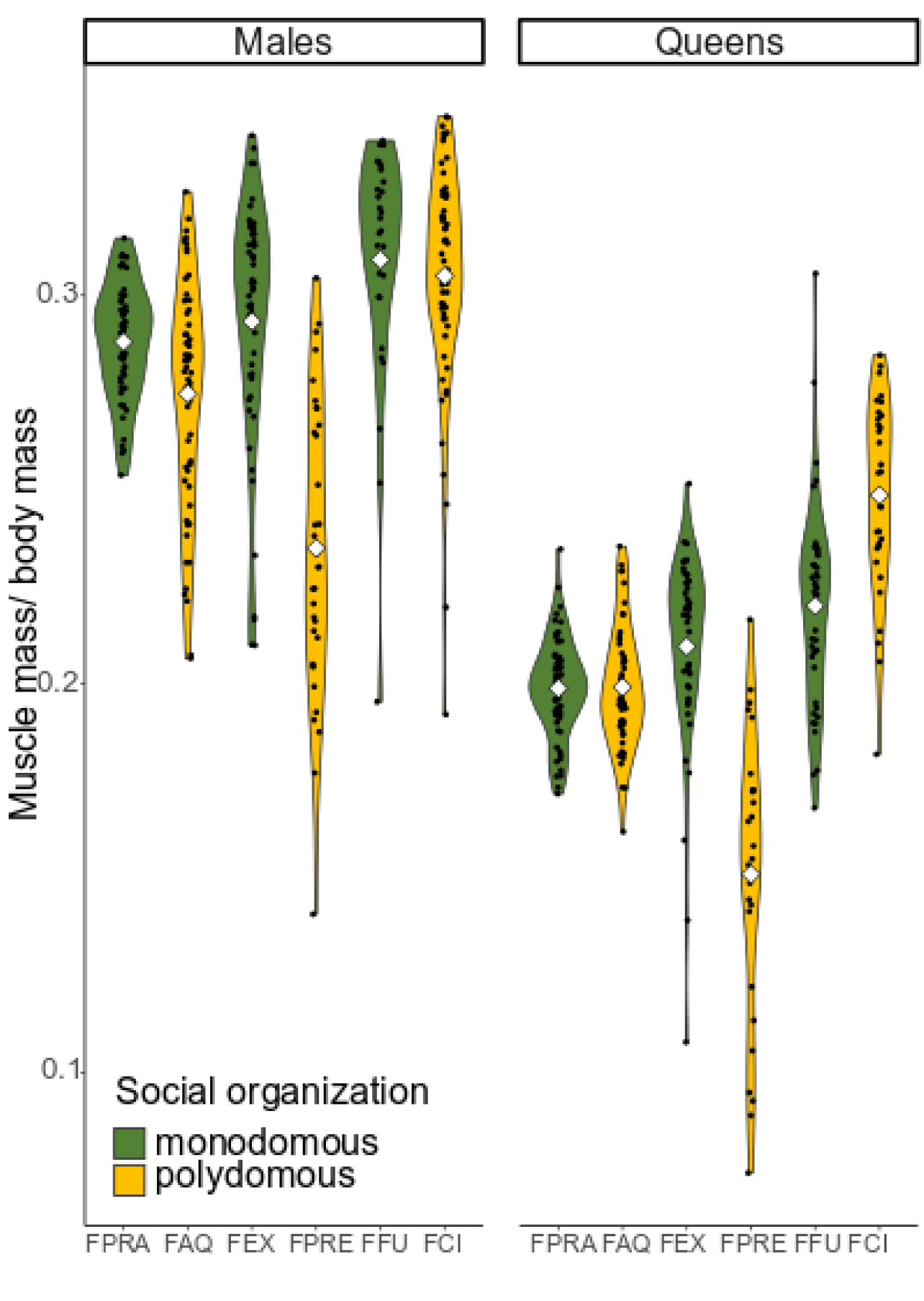
Flight muscle mass to body mass ratio by species and by sex. All data are visualized as data points and density plots, the mean is visualized with a diamond shape. The species are abbreviated as follows: FPRA = *F. pratensis*, FAQ = *F. aquilonia*, FEX = *F.exsecta*, FPRE = *F. pressilabris*, FFU = *F. fusca*, FCI = *F.cinerea*. The males have significantly larger muscle ratios than the queens, and the monodomous males have larger muscle ratios than the polydomous males (Table 2).

**Table 2:** Analyses of the body proportions and biochemical measurements. In all the analyses, the social organization (monodomous/polydomous), sex, and their interaction were used as fixed effects, except for the triglyceride measurements where there are data only for queens and thus only the social organization is used as fixed effect. Species and nest were used as random effects. The Benjamini-Hochberg procedure (1995) was used to correct the parameter p-values to decrease the false discovery rate (FDR). Tests where the p-value is smaller than the FDR critical value are considered significant. Non-significant results are presented in grey.

| Response | robust LMM |  |  | LMM |  |  |  |  |
| --- | --- | --- | --- | --- | --- | --- | --- | --- |
| Parameter names |  |  |  |  |  |  |  |  |
| Total body mass | $\beta$ | SE | t-value | $\beta$ | SE | t-value | p | BH critical value |
| Intercept (Monodomous males) | 0.018 | 0.007 | 2.520 | 1.803e-02 | 6.095e-03 | 2.959 | 0.0415 |  |
| Polydomous | -0.006 | 0.010 | -0.564 | -5.618e-03 | 8.619e-03 | -0.652 | 0.5501 | 0.615 |
| Queens | 0.006 | 0.000 | 24.620 | 6.845e-03 | 3.157e-04 | 21.685 | <2e-16 | <0.001 |
| Interaction | 0.000 | 0.000 | -0.721 | -8.724e-04 | 4.393e-04 | -1.986 | 0.0473 | 0.112 |
| Abdomen ratio |  |  |  |  |  |  |  |  |
| Intercept (Monodomous males) | 0.405 | 0.030 | 13.562 | 0.408 | 0.026 | 15.848 | 0.000 |  |
| Polydomous | 0.003 | 0.042 | 0.066 | 0.001 | 0.036 | 0.027 | 0.980 | 0.980 |
| Queens | 0.014 | 0.003 | 4.258 | 0.015 | 0.004 | 4.116 | 0.000 | <0.001 |
| Interaction | 0.007 | 0.005 | 1.485 | 0.004 | 0.005 | 0.723 | 0.470 | 0.558 |
| Thorax ratio |  |  |  |  |  |  |  |  |
| Intercept (Monodomous males) | 0.523 | 0.012 | 42.960 | 0.521 | 0.011 | 48.475 | 0.000 |  |
| Polydomous | -0.016 | 0.017 | -0.940 | -0.016 | 0.015 | -1.036 | 0.357 | 0.522 |
| Queens | -0.074 | 0.003 | -26.840 | -0.075 | 0.003 | -24.373 | <2e-16 | <0.001 |
| Interaction | 0.000 | 0.004 | 0.120 | 0.003 | 0.004 | 0.768 | 0.442 | 0.558 |
| Muscle ratio |  |  |  |  |  |  |  |  |
| Intercept (Monodomous males) | 0.298 | 0.018 | 16.538 | 0.295 | 0.017 | 17.042 | 5.93e-05 |  |
| Polydomous | -0.027 | 0.025 | -1.043 | -0.026 | 0.024 | -1.052 | 0.351 | 0.522 |
| Queens | -0.087 | 0.003 | -26.902 | -0.085 | 0.004 | -22.374 | <2e-16 | <0.001 |
| Interaction | 0.019 | 0.005 | 4.182 | 0.019 | 0.005 | 3.424 | 0.001 | 0.002 |
| Protein concentration |  |  |  |  |  |  |  |  |
| Intercept (Monodomous males) | 3.074 | 0.231 | 13.307 | 3.269 | 0.264 | 12.370 | 1.32e-05 |  |
| Polydomous | 0.699 | 0.324 | 2.159 | 0.742 | 0.370 | 2.004 | 0.092 | 0.178 |
| Queens | -1.316 | 0.204 | -6.469 | -1.448 | 0.251 | -5.769 | 1.94e-08 | <0.001 |
| Interaction | -0.128 | 0.292 | -0.440 | -0.037 | 0.361 | -0.103 | 0.918 | 0.969 |
| Glycogen concentration |  |  |  |  |  |  |  |  |
| Intercept (Monodomous males) | 1.120 | 0.259 | 4.320 | 1.243 | 0.235 | 5.297 | 0.003 |  |
| Polydomous | -0.387 | 0.365 | -1.059 | -0.468 | 0.330 | -1.418 | 0.218 | 0.376 |
| Queens | 0.189 | 0.125 | 1.512 | 0.258 | 0.154 | 1.679 | 0.094 | 0.178 |
| Interaction | 0.478 | 0.179 | 2.664 | 0.480 | 0.221 | 2.176 | 0.030 | 0.082 |
| Triglyceride concentration (queens) |  |  |  |  |  |  |  |  |
| Intercept (Monodomous) | 0.382 | 0.124 | 3.092 | 0.442 | 0.122 | 3.631 | 0.0265 |  |
| Polydomous | -0.113 | 0.174 | -0.650 | -0.153 | 0.172 | -0.888 | 0.430 | 0.558 |

The queens were overall larger than males of the same species (Table 2, Appendix 2: Figure 1). There was a lot of size variation among the species, but it was not consistently explained by the social organization (Table 2, Appendix 2: Figure 1). The body ratios were opposite in males and queens: the queens had larger abdomen ratios than males, whereas males had larger thorax ratios than queens (Table 2). *Coptoformica* queens had larger heads than other *Formica* queens, and their other body ratios were accordingly affected: especially the abdomens were smaller than in other *Formica* (Appendix 2: Figure 1).

### MICROSCOPIC MUSCLE STRUCTURE

Overall, the muscle structure analysis revealed that all species had well-structured flight muscles with rather evenly-organised myofibrils (average CV of distances per image for each species and sex varies between 0.218-0.257 with small standard deviations, Appendix 1: Table 1). All the species studied had large quantities of mitochondria almost filling the spaces among the myofibrils (on average, 40.0% of the total cross section area), though individual mitochondria profile areas were small and their shapes were closer to round instead of elongated and tubular (Figure 2, Appendix 1: Table 1). There was large variation within species, but only small variation among species (Appendix 3: Figures 1 and 2). There were signs of minor degradation of mitochondria, but it was not extensive (Appendix 3: Figures 3 and 4). None of the differences in muscle structures could be explained by social organization or sex (Appendix 3: Figures 1 and 2 and Table 1).

**Figure 2:**
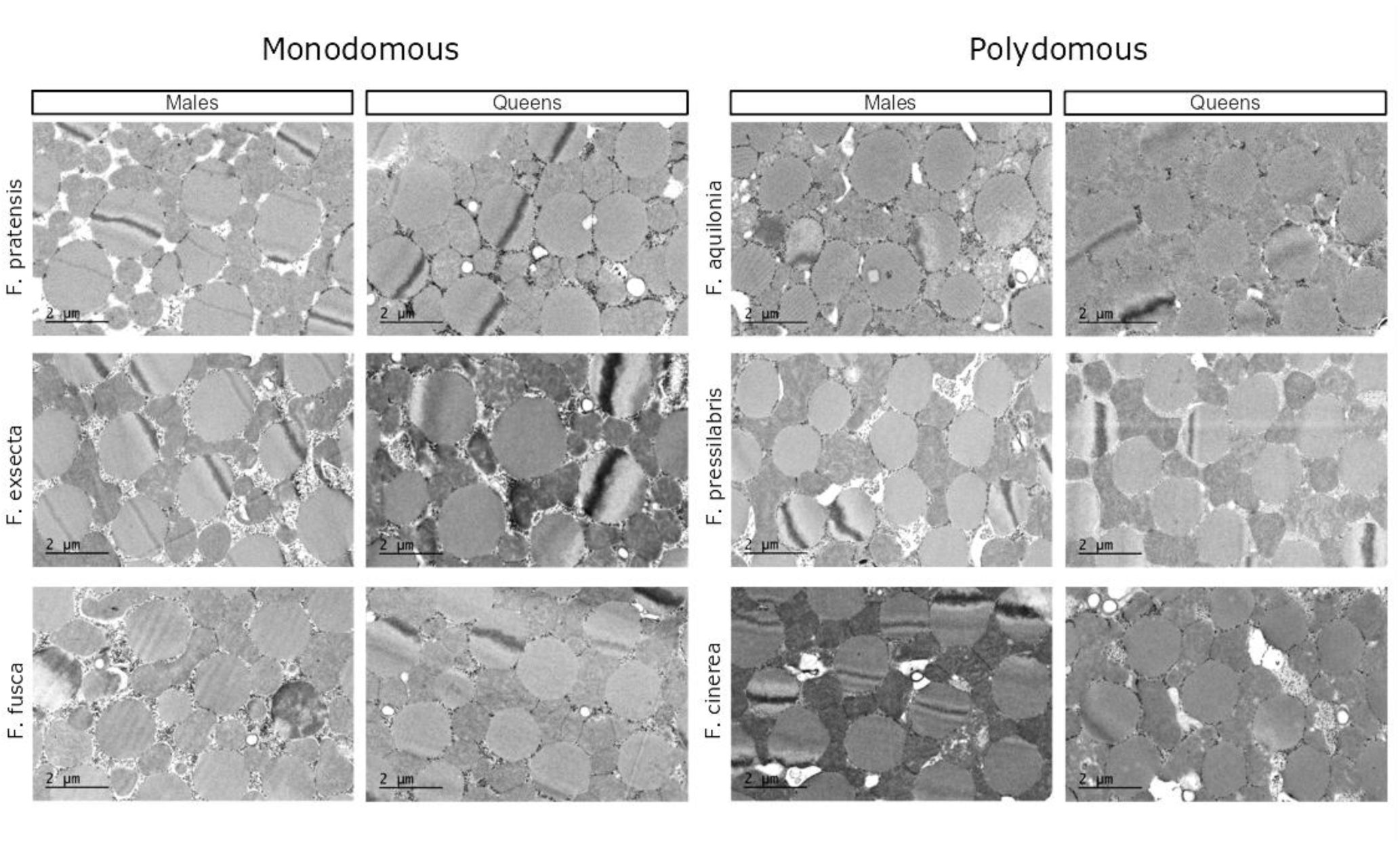
Typical transmission electron microscopy images of flight muscle tissue for all of the sampled *Formica* species and both sexes. All species have functioning muscles, and there are no statistically significant differences in the mitochondria and myofibril sizes or structures among the species. Myofibril cross sections are visible as round, equally distributed shapes in the images. Mitochondria are the unevenly distributed oval shapes, with cristae visible inside. The black granules between the cell organelles are glycogen.

**Figure 3:**
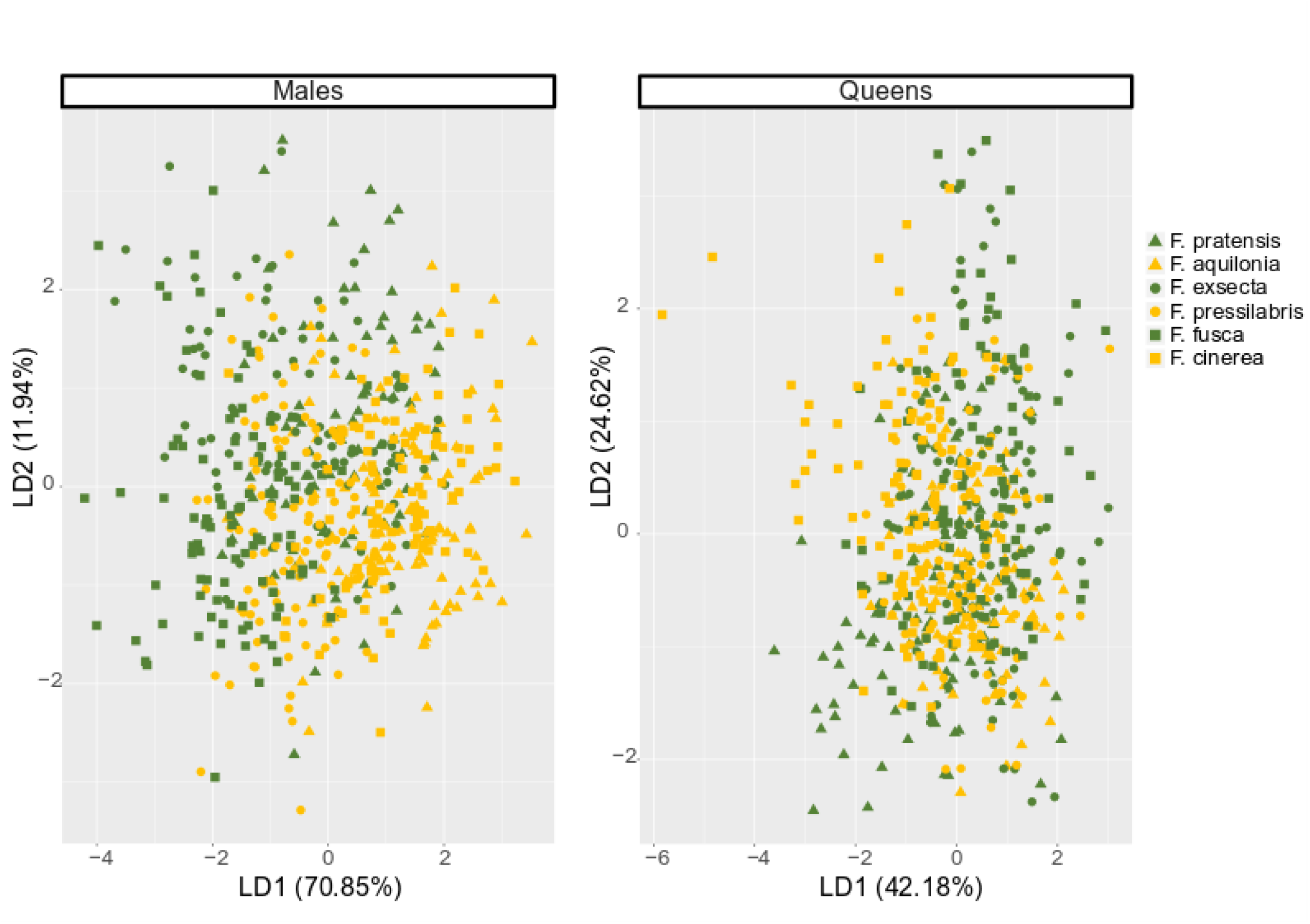
Linear discriminant analyses for each species by microscopical muscle structure data. The symbol denotes the subgenus: triangle = *Formica rufa* –group, circle= *Coptoformica*, rectangle: *Serviformica*. The color denotes social organization: dark green = monodomous, yellow = polydomous.

**Figure 4:**
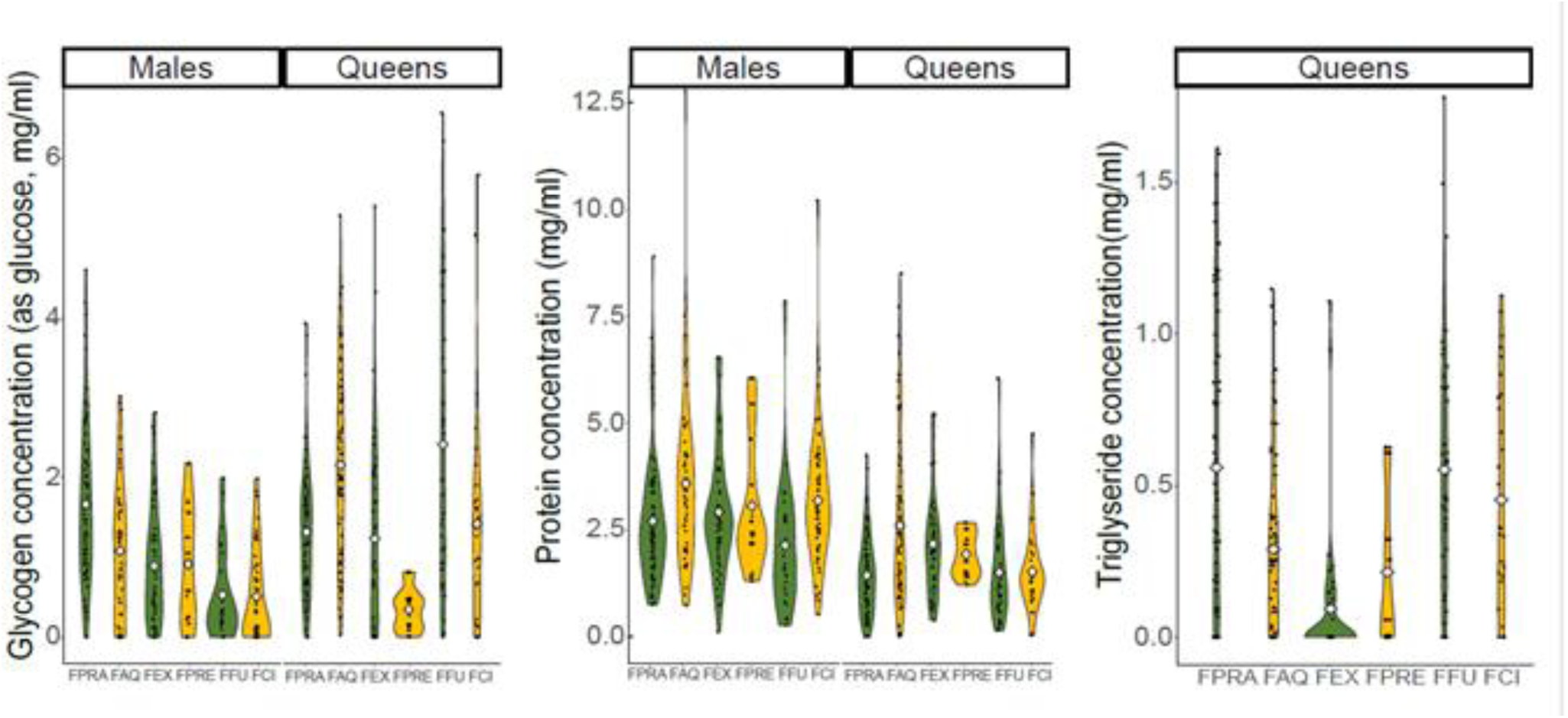
Mass-corrected biochemical resources by species and by sex. All data are visualized as data points and density plots, the mean is visualized with a diamond shape. The species are abbreviated as follows: FPRA = *F. pratensis*, FAQ = *F. aquilonia*, FEX = *F.exsecta*, FPRE = *F. pressilabris*, FFU = *F. fusca*, FCI = *F.cinerea*. The social organization of each species is highlighted with color: dark green = monodomous, yellow = polydomous. Social organization does not explain any of the variation (Table 2). a) Glycogen concentration, b) protein concentration, c) triglyceride concentration for the queens.

For the muscle structure data, the sex-specific LDA models assigned the samples to two groups based on social organization with 63.8% (kappa=0.28) and 73.7% (kappa=0.47) accuracy for queens and males, respectively. Even though the single measurements do not show differentiation, and the LDA accuracy overall is not high, this still suggests there is minor differentiation in overall muscle structures between the two social organizations. The LDA models were not much better able to find species-level differentiation based on muscle structures, with model accuracies of 40.1% (kappa=0.28) for the queens and 50.7% (kappa=0.41) for the males. (Figure 3)

### GLYCOGEN, TRIGLYCERIDE AND PROTEIN ANALYSIS

There were species-specific differences in the amounts of metabolic resources, but none of this variation was consistently explained by the social organization (Table 2). Glycogen concentrations in males were more similar in closely related species, whereas queens had large variation among the species regardless of the phylogeny (Figure 4a). The queens also had greater overall variation in glycogen concentration (Appendix 2: Table 1).

The mean soluble protein concentrations were similar in the six species, with no effects of the social organization (Figure 4b, Table 2). Males had more protein than queens and the variation was larger in males than in queens (Figure 4b, Table 2, Appendix 2: Table 1).

Males did not have measurable amounts of stored triglyceride – only 16.2% of the males in our dataset had values higher than zero, and some of these were among the pooled *F. pressilabris* individuals, making the actual number likely even smaller. As it is likely that these rare positive values reflect their last meal rather than stored resources, the male triglyceride measurements were not analyzed further. Additionally, some of the *Coptoformica* queens had such low amounts of triglyceride that it could not be measured with our assay. Overall, the queen triglyceride concentration varied greatly among species (Figure 4c).

### OVERALL DIFFERENTIATION

In males, there was a positive correlation between individual size and muscle ratio: larger individuals also had proportionally larger muscles. Similar, though not statistically significant, trend was seen in queens. The thorax ratio and abdomen ratio were strongly negatively correlated in both sexes. Interestingly, thorax ratio and muscle ratio did not correlate in queens, although such correlation was assumed in previous studies. They did correlate positively in males. (Appendix 2: Figure 2)

For the body ratio and biochemical data, the sex-specific LDA models assigned the samples to groups based on social organization with 55.3% (kappa=0.10) and 81.3% (kappa=0.62) accuracy for queens and males, respectively, and to groups based on species with 74.7% (kappa= 0.68) and 86.7% (kappa=0.84) accuracy for queens and males, respectively. Thus, these models were better able to separate both groupings for the male data. Overall clustering reflects more the subgenus than the social organization (Figure 5).

**Figure 5:**
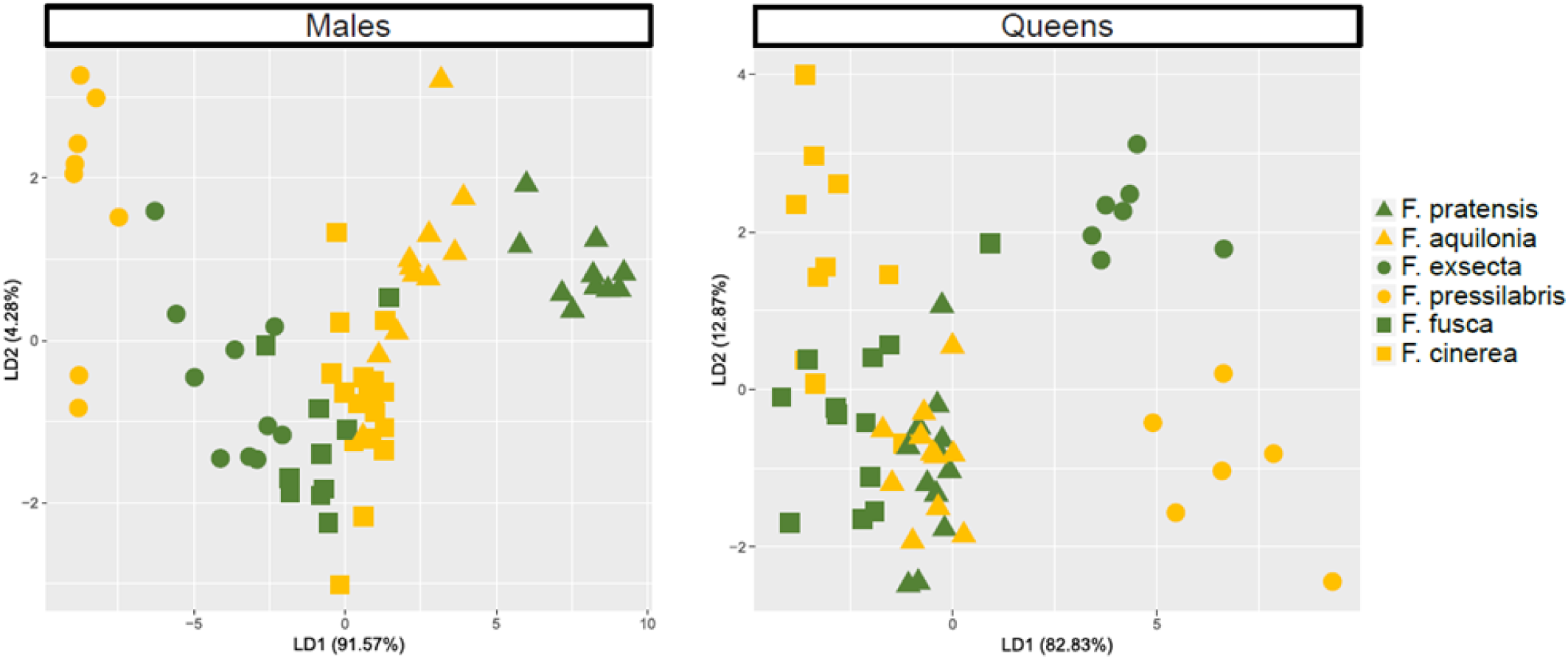
Linear discriminant analyses for each species by morphological and biochemical data. The symbol denotes the subgenus: triangle = *Formica rufa* –group, circle = *Coptoformica*, rectangle = *Serviformica*. The color denotes social organization: dark green = monodomous, yellow = polydomous.

### WITHIN-SPECIES VARIATION

*F. exsecta* had smaller individuals in the polydomous than in the monodomous populations, whereas *F. pressilabris* had no size difference between the social organizations – the individuals of this species were always very small. In *F. pressilabris*, the queens were smaller than the males, which differs from the general pattern in *Formica* and ants in general. In *F. exsecta*, both males and queens had smaller muscle ratio in the polydomous population than in the monodomous populations, whereas in *F. pressilabris* there was no effect of social organization on the muscle ratio. There were no intraspecific differences in the biochemical measurements between the two social organizations. (Figure 6, Appendix 1: Table 2, Appendix 2: Table 2).

**Figure 6:**
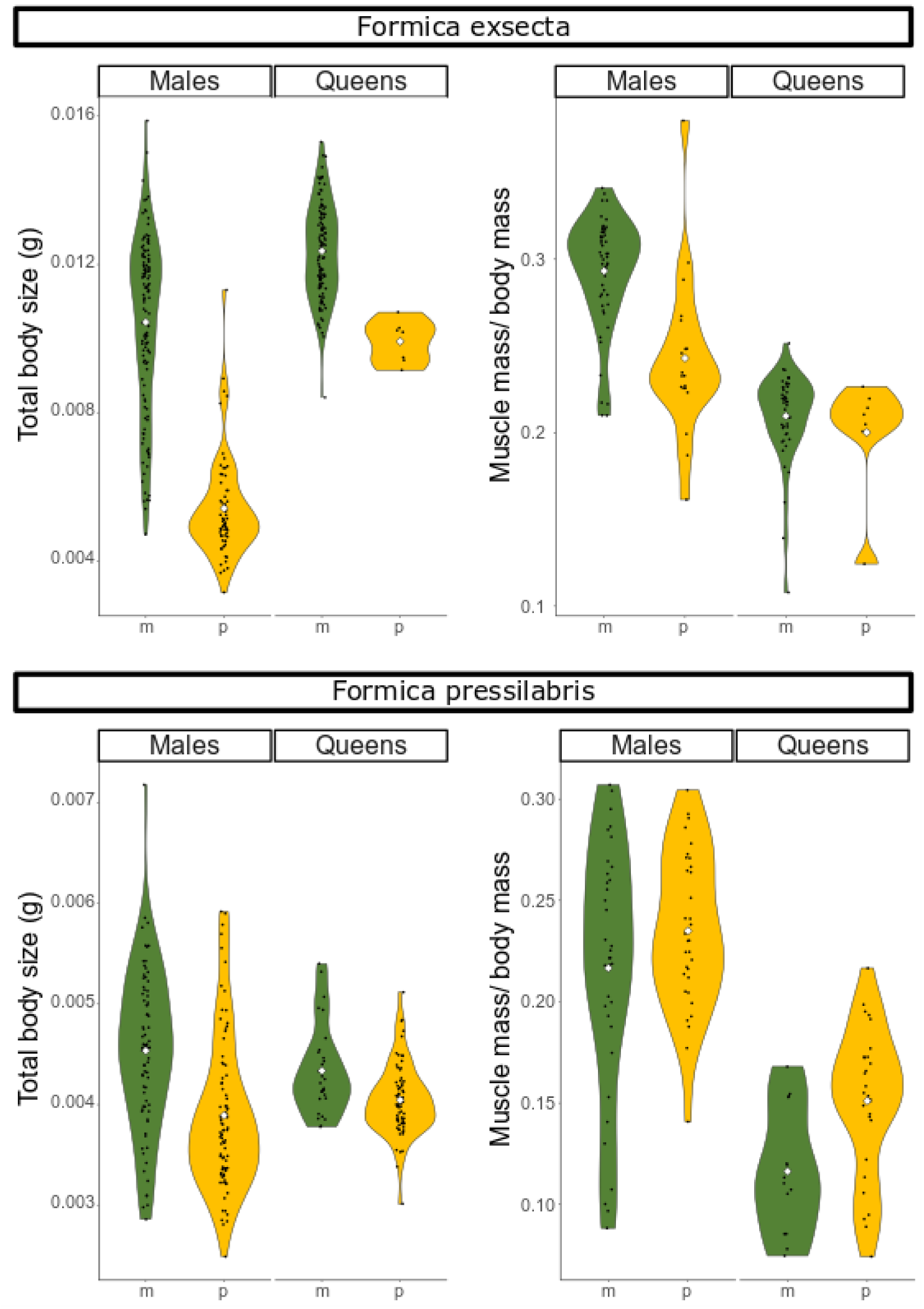
Total body size and muscle mass ratio by social organization and by sex for *F. exsecta* and *F. pressilabris*. All data are visualized as density plots with all data points, the mean is visualized with a diamond shape. The color denotes social organization: dark green = monodomous (m), yellow = polydomous (p). The differences between the two social organizations are significant in *F. exsecta* but not in *F. pressilabris* (Appendix 2: Table 2).

## DISCUSSION

Our hypothesis that the philopatric queens of the polydomous *Formica* species would have smaller resource allocation to dispersal traits than queens of the monodomous species was not met. Thus, social polymorphism in *Formica* ants is a result of queen behavior rather than dispersal ability. However, male muscle ratios are smaller in the polydomous species, showing strong coevolution of the sexes: male morphology is slightly differentiated between the queen-behavior-driven social organizations. The hypothesis for sex-biased dispersal was met, as males have larger flight muscle ratios than the queens across the genus.

### *FORMICA* FLIGHT ABILITY

In general, *Formica* muscle sizes are adequate for flight in all the studied species. Marden (1987) specifies a threshold of 12-16% of minimum muscle ratio for successful take off, and all of our species exceed this. The only exception is a proportion of *F*. *pressilabris* queens that have smaller muscle ratios than this, suggesting that in this species the queen flight ability is partially compromised. *Formica pressilabris* seems to be overall the poorest disperser in our dataset, with by far the smallest muscle ratios in both sexes, and the lowest glycogen concentrations in the queens.

In addition to muscle ratios, also the muscle structures indicate functional muscles in all the species and both sexes. The total mitochondrial area, 40% of the muscle cross sections, compares well to other flying insects. *Libellula* dragonflies, good examples of strong flyers because they spend a third of their time in flight (Marden et al. 1996), have only slightly larger mitochondria areas (46%, Marden 1989). Even *F. pressilabris*, with its smallest muscles, has good muscle structures, and likely flies adequately when the muscles are large enough. However, ant muscle structures do differ from more actively flying insects: insect flight muscles often have large fused mitochondria that tightly envelope the myofibrils (Edwards and Ruska 1955, Sohal and Allison 1971, Sohal et al. 1972), whereas most of the mitochondria in our ant samples are singular and round.

Of our study species, the dispersal of *F. exsecta* has been shown to happen mostly over short distances (Sundström et al. 2003, Vitikainen et al. 2015). *Formica exsecta* has rather average dispersal traits when compared to the other species in our dataset, suggesting that none of the species are exceptionally good dispersers. This is in line with studies on other ants. For example *Solenopsis invicta* queens seem to have resources for an average flight of only 45 minutes, thus dispersing long distances only passively by wind (Vogt et al. 2000).

### SEX BIASED FLIGHT ABILITY

The muscle ratios are larger in males, indicating that *Formica* males are stronger flyers than *Formica* queens, regardless of the social organization, concurring with the male-biased dispersal shown in gene flow analyses (Sundström et al. 2005). Interestingly though, glycogen concentration does not have overall sex differences. It is possible that the queen and male flight has different energetic needs because of larger abdomen drag in queens (Helms 2018). In *S. invicta,* queens increase their metabolic rate more than males during flight (Vogt et al. 2000). Thus glycogen resources cannot be directly compared between sexes.

In *F. fusca*, gene flow was shown to be unbiased between the sexes, or even slightly female biased (Johansson et al. 2018), in contrast to the general male bias in *Formica* ants (Sundström et al. 2005). In our data the males of this species still do have larger muscles than the queens. However, the glycogen reserves of *F. fusca* males are low compared to other *Formica* males, and this may limit their flight duration. Observational studies on dispersal and mating behavior in natural populations have not been done for *F. fusca*, but Hannonen et al. (2004) suggest both male and queen dispersal may be more limited especially in older populations. In a related and ecologically similar *F. selysi*, both sexes appear in the mating swarms close to the natal colonies, but also here their flight after mating was not monitored (Fontcuberta et al. 2021).

### THE EFFECT OF SOCIAL ORGANIZATION

Against our main hypothesis, monodomous queens do not have clearly higher allocation to dispersal traits than the polydomous queens. Dispersal abilities vary among species, showing the importance of species specific ecological and evolutionary constraints in dispersal evolution, but there is no overall pattern due to the social organization. Thus, queen philopatry in polydomous societies is likely a behavioral decision and not caused by queen morphology. It is still possible that we have missed the most philopatric queens in our sampling, if they are rare or do not come to the nest surface even for mating – but this seems unlikely. For example, in *F. pressilabris,* also the queens with so small flight muscle ratios that they are likely unable to fly, still appeared on the nest surface where we sampled them.

In our male data, the muscle ratios are overall slightly larger in monodomous than in polydomous males, indicating that flight ability is lower in polydomous males. Although this effect is small, it is biologically meaningful. Muscle ratio is known to straightforwardly affect the flight ability of the individuals (Marden 1987). Additionally, social organization affects the variance of the body proportions, including muscle ratio that varies more in the polydomous species. This indicates that selection on mating and breeding strategies are disruptive in polydomous societies and, as a result, they have more variation in dispersal traits compared to monodomous societies. If there is simultaneous selection for dispersal and philopatry in the queens, it is possible that the same is true for the males: if local mating with the philopatric queens is possible, a proportion of the males may be selected not to allocate their resources to flight muscles. Due to low relatedness in polydomous societies, the risk of inbreeding is smaller in them than in monodomous societies, which makes evolution of increased philopatry of both sexes more likely (Vitikainen et al. 2015, Hakala et al. 2019).

The combined microscopic muscle structures are slightly differentiated between the two social organizations based on LDA, with less separation by species. It is not clear how this is reflected in the muscle function, but the result supports our conclusion that although the evolutionary changes in dispersal traits in connection to social organization are small in *Formica* ants, they do exist.

### COEVOLUTION OF DISPERSAL AND MATING

When considering the possibility of increased philopatry in *Formica* males, we need to consider their mating system. *Formica* queens can use pheromones to attract males to mating swarms. Males can also visually locate male aggregations that form when several males locate the same queens. Instead of mating away from their natal colony, queens can use their pheromones to attract males towards their natal colony – and likely do so in all the polygynous and polydomous species where daughter queens are recruited back. However, the amount of studies analyzing mating behavior in *Formica* is very small and with a small number of species – from our study species only *F. aquilonia* and *F. exsecta* have been studied in this respect. (Kannowski and Johnson 1969, Cherix et al. 1991, 1993, Walter et al. 1993, Fortelius 2005, Martin et al. 2014, Fontcuberta et al. 2021).

In polydomous ants, there are plenty of opportunities for local mating, and evolution of philopatric males seems possible in the light of our data. In supercolonial *F. paralugubris,* mating with nestmates is common based on genetic data (Chapuisat et al. 1997, Chapuisat and Keller 1999). Although there is not enough direct data on male behavior, we hypothesize that it has changed in the polydomous species due to selection for local mating with the philopatric queens. As an extreme example, simultaneous selection for both local mating and dispersal leads to evolution of separate morphs in wing-dimorphic insects, due to a trade-off between sperm amount and flight (Zera and Denno 1997, Saglam et al. 2008). In ants such male dimorphism is very rare, but does exist in *Cardiocondyla* ants (Kinomura and Yamauchi 1987, Heinze and Hölldobler 1993). We do not have data on male sperm amounts, but their total protein amount can be considered to somewhat represent the amount of seminal fluid, which contains large amounts of proteins (Avila et al. 2011). The concentrations vary substantially within species, showing that there are possibilities for selection. However, we found no correlation between muscle size and protein concentration in males, and no differences between the two social organizations, revealing no clear division to two different male types in general. However, in *F. exsecta*, there are two male size morphs (Fortelius et al. 1987) that we will discuss in a later section.

Proper analysis of both sperm quantity and mating behavior across the different social organizations would be interesting. Polydomous *Formica* is substantially easier to induce to mate in the laboratory (Fortelius 1987, but see Martin et al. 2014), which indicates that in polydomous species there indeed has been evolution towards local mating, possibly through decoupling their mating propensity from flight.

### DIFFERING ECOLOGIES OF THE STUDY SPECIES

Males can be more accurately assigned to both species and social organization than queens with the LDA based on combined muscle ratio, body proportion and biochemical data. These results indicate that there may be slightly more diversification of male dispersal traits, possibly because males are under smaller evolutionary constraints and trade-offs than the queens and thus more free to evolve (Helms and Kaspari 2015, Hakala et al. 2019). One such evolutionary constraint for the queens is the distinct head shape of *Coptoformica*: they have long heads with strong jaw muscles because workers chop grass into nest-building material, and the queens share this morphological feature with them (Seifert 2000). This likely restricts their other body proportions and thus their dispersal evolution – resulting in smaller abdomen ratios and smaller triglyceride amounts, compared to other *Formica* queens.

The most obvious ecological difference affecting dispersal evolution in our study species is the mode of nest founding: Only the two *Serviformica* species are capable of independent nest founding, and of them only *F. fusca* is an obligate independent founder whereas *F. cinerea* can also use dependent founding (Collingwood 1979, Goropashnaya et al. 2001). The other study species are dependently founding or temporary social parasites that sneak into the nests of other *Formica* ants instead of founding their own nests (Collingwood 1979, Buschinger 2009). Following the logic of the found or fly hypothesis, which states that there is a trade-off between flight and nest founding abilities (Helms and Kaspari 2014, 2015), an obligatory independent founder should have higher nest founding resources and thus poorer flight resources compared to species that have other options for nest founding. This is the case in fire ants where the leaner queens with proportionally larger muscles are able to fly longer, but act like parasites due to smaller nest founding resources (Helms and Godfrey 2016). Based on our data, the found or fly trade-off does not play out as expected in *Formica*: the socially parasitic queens do not have larger muscle ratios or consistently poorer nest founding resources. On the contrary, the highest queen muscle ratios are in *Serviformica* species, and the highest triglyceride concentrations are in one of the social parasites, *F. pratensis*. The only obligate independent nest founder, *F. fusca*, has better queen resources for both flying and nest founding than average *Formica*. The pioneer ecology of *F. fusca* (Hannonen et al. 2004, Johansson et al. 2018) is likely to create strong selection pressure for both good dispersal ability and good nest founding ability, whereas the parasitic species may be under other competing selection pressures arising from their parasitic lifestyles (Brandt et al. 2005, Hakala et al. 2019).

Our study species are very differently sized, and thus we mostly discuss their dispersal resources as size-corrected ratios and concentrations. However, the relationships between actual body size, dispersal ability and colony resource allocation are very complex. Because metabolism scales with body size in such a way that larger individuals have relatively smaller metabolic rates, the energetic costs of all activities, including flight and maintenance of flight muscles, depend on the body size (Chown and Nicolson 2004, Suarez et al. 2004, Chown et al. 2007). Thus, absolute body size plays a role in dispersal ability. According to our results, the body size correlates positively with some of the size-corrected measurements, which indicates that larger individuals are overall more able to have good dispersal resources. Direct measurements of dispersal physiology and metabolic rates would aid in confirming this hypothesis.

Smaller queens have long been associated with the more complex ant societies, a phenomenon connected to the so-called polygyny syndrome, and ultimately limited dispersal (Rosengren and Pamilo 1983, Keller 1993). Producing smaller sexuals is a sign of smaller colony level resource allocation. According to our results the body size is not consistently explained by social organization at the species level: the *F. rufa* group species and *Coptoformica* species follow this pattern, whereas the *Serviformica* species do not.

### WITHIN-SPECIES VARIATION

*F. exsecta* males are size dimorphic: tiny males are found in predominantly polygynous and polydomous populations, and large males in monogynous and monodomous populations (Fortelius et al. 1987). Monogynous populations have rather large male size variation (Vitikainen et al. 2015), and it is easy to hypothesize that all populations have the potential to produce small males. However, for some reason small males are selected for in the polydomous populations, leading to reduction or loss of big males in them. Previously smaller males have been considered better dispersers (Fortelius et al. 1987), but in our data small male size correlates with substantially lower muscle mass / body mass ratios and thus likely with poorer dispersal. More work on dispersal behavior of *F. exsecta* males is needed to resolve this contradiction. In addition to males, also *F. exsecta* queens are smaller in the polydomous compared to monodomous population*s*. The evolution of decreased size of sexuals in polydomous *F. exsecta* might reflect the same selection pressures that have led to the evolution of tiny sexual castes in the closely related *F. pressilabris*.

Interestingly, although both *F. exsecta* and *F. pressilabris* are socially polymorphic, only *F. exsecta* has a size difference between the two social organizations. Additionally, in *F. exsecta* but not in *F. pressilabris*, muscle ratios are smaller in the polydomous population in both sexes. This difference reflects the overall mismatch between dispersal behavior and dispersal ability that we see in the interspecific comparisons.

Trait variation is the currency evolution uses – not just genetic variation but to some extent also plastic variation (Pfennig et al. 2010, Moczek et al. 2011). In our study, the within-species variation of most of the measured traits is large. This could lead to condition dependent dispersal decisions at the individual level (Bowler and Benton 2004), as it seems to do in *F. truncorum* (Sundström 1995). We suggest that switch from monodomous to polydomous social organization likely starts as a plastic response to environmental pressure and individual condition. Polydomous populations may subsequently lose some of their original plasticity and variation, leading to more fixed philopatric traits. This could explain the evolution of small sexuals in commonly polydomous *F. pressilabris*, although the exact selection pressures should be further analyzed.

It is intriguing that although the variation in dispersal traits is large and does to some extent correlate with social organization in our data, the patterns within-species in *Coptoformica* and between species across the whole genus are somewhat contradictory. There are clearly additional forces or restrictions of selection at play. One of these is undoubtedly the *Formica* supergene controlling queen number, carried at least in fragmented form in most of our study species, though not all have been analyzed (Purcell et al. 2014, Brelsford et al. 2020, Lagunas-Roblesa et al. 2024, Sigeman et al. 2024). In *F. selysi*, the model species for the supergene, there are colony-level effects in the numbers and size of sexuals produced between the two supergene alleles, and the queens carrying the allele allowing polygyny appear less at the mating swarms (Fontcuberta et al. 2021). Similar supergenes controlling queen numbers have evolved in many unrelated ant taxa (Kay et al. 2022), which suggests similar selection pressures govern their evolution. Dispersal is potentially a major one. Once a supergene exists, it is a strong driver and restrictor of further evolution (Chapuisat 2023), and finding the was the supergene connects to the functional mechanisms of controlling the colony queen number (morphology, physiology, behavior of workers and/or sexuals) is a relevant future direction. The curios disappearance of the polygyny-associated allele in the obligatorily polygynous and polydomous species also begs for further studies, which would aid the discussion of our results (Lagunas-Roblesa et al. 2024, Sigeman et al. 2024)

### POSSIBILITY OF DISPERSAL CONFLICT

As the supercolonial *Formica* queens are able to fly but are still philopatric, their societies seem to waste substantial amounts of resources when producing them. The young queens are provided with functional flight muscles but many of them do not use them. We suggest the high levels of philopatry could be interpreted as selfish behavior of the young queens, to avoid the risks of dispersal and to start reproducing sooner without independent colony founding. Selfish behavior is hypothesized to increase in supercolonial ant societies (Helanterä et al. 2009) due to low relatedness among nestmates. It indeed seems that the supercolonial *Formica* societies have more queens than they need, even hundreds per nest and thousands per polydomous colony (Stockan and Robinson 2016). If excessive amounts of queens stay in their natal nest, their behavior can even be seen as parasitic (Rosengren et al. 1993a).

We further suggest that increased selfish philopatry of the queens may lead to conflict over dispersal in the supercolonial societies, where the other members of the society would prefer the young queens to disperse more than they are willing to (Motro 1983, Starrfelt and Kokko 2010, Hakala et al. 2019). Previous research has suggested that the polygynous (and polydomous) populations of *Formica* have density dependent selection for both dispersing and philopatric queens (Rosengren et al. 1993). However, to our knowledge this theory has not been rigorously tested, or even mathematically analyzed for complex haplodiploid societies, and no density dependence in their dispersal has been shown in natural populations. Instead, we suggest it is possible that in the extremely polygynous and polydomous *Formica* populations there is a feedback loop between philopatry leading to low relatedness that leads to more selfish queens that are even more philopatric (Hakala et al. 2019) – altering the evolutionary trajectory of their societies, and even the survival of the species in todays fragmented landscapes (Hakala et al. 2023).

## CONCLUSIONS

All *Formica* species in our study have adequate dispersal resources, even though queens in half of our study species are highly philopatric. Overall dispersal does not correlate well with morphological proxies for dispersal ability, but it is more of a behavioral trait. Similarly it has been shown in the Mediterranean fruit fly that a measure of flight performance (vertical flight force output) did not differ between individuals classified as dispersers vs. sedentary, whereas the number and duration of voluntary flight bursts did (Steyn et al. 2016). Together with our results, this suggests that the ability to disperse may not be the main driver of dispersal events in nature, but behavior has a larger role than traditionally appreciated. Evolution can be driven by behavior, followed by morphological changes.

We suggest that the polydomous and supercolonial societies in *Formica* evolve through a change in queen dispersal behavior, independent of their morphology. Interestingly, the observed limitation in polydomous male dispersal morphology reveals increased philopatry in polydomous males, too. Male dispersal ability evolving as a response to queen-behavior-created social organization is a novel example of sexual coevolution in ants that are usually studied for their social female traits alone.

## ACKNOWLEDGEMENTS

Heini Ali-Kovero, Hanne Henriksson and Kirsi Kähkönen from the Molecular Ecology and Systematic laboratory and Mervi Lindman and Arja Strandell from the Electron Microscopy Unit offered invaluable assistance. Mari Teesalu helped with the biochemical protocols. The personnel of the Tvärminne Zoological station assisted with field permits and other aspects of field work. Matti Leponiemi, Tanya Troitsky and Eeva Vakkari helped with field work and handling the specimens. Dimitri Stucki gave valuable comments about the statistical analyses. Heike Feldhaar, Kristjan Niitepõld and Mark Brown gave excellent comments during the examination of SH:s PhD thesis that included an earlier version of this manuscript. Our work was funded by the Academy of Finland (#140990, #135970, #251337 and #284666). Additionally, SH was funded by Finnish Cultural Foundation and Alfred Kordelin Foundation, and supported by Swiss National Science Foundation grant (#310030-207642 to Michel Chapuisat) during the last phases of the work. HH was funded by Kone foundation and EJ and IB were funded by Biocenter Finland.

