## Appendix 2 for "DISPERSAL BEHAVIOR RATHER THAN DISPERSAL MORPHOLOGY CREATES SOCIAL POLYMORPHISM IN FORMICA ANTS"

### APPENDIX 2: BODY PROPORTIONS AND BIOCHEMICAL RESOURCES

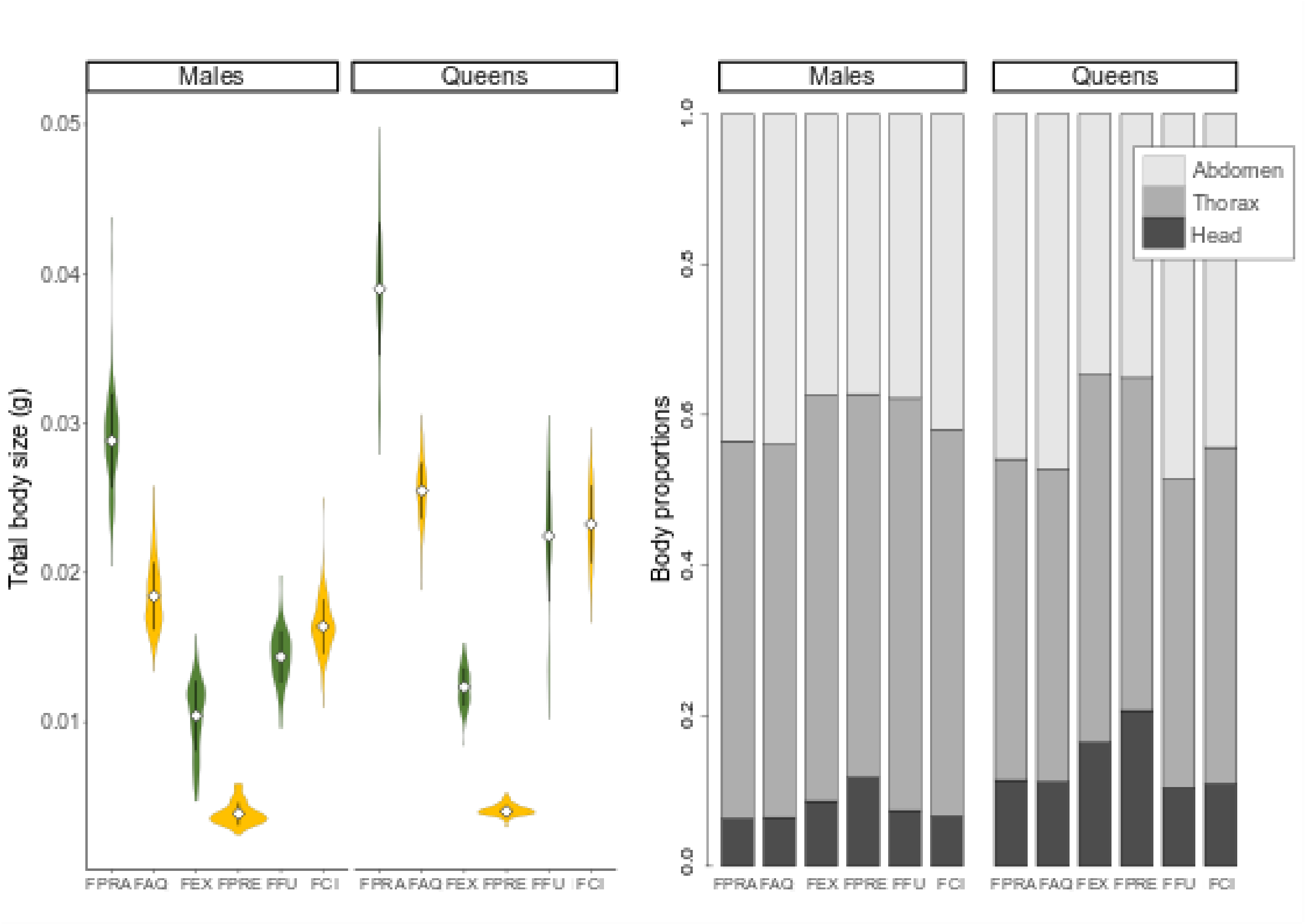


**Figure 1. Left panel: Body mass by species and by sex.** All data are visualized as density plots (violin plots), the mean is visualized with a diamond shape and +- standard deviation with whiskers. The species are abbreviated as follows: FPRA = *F. pratensis*, FAQ = *F. aquilonia*, FEX = *F.exsecta*, FPRE = *F. pressilabris*, FFU = *F. fusca*, FCI = *F.cinerea*. The social organization of each species is highlighted with color: dark green = monodomous, yellow = polydomous. The data of *F. exsecta* and *F. pressilabris* used for this figure include only the main populations that have the typical social organization. For within-species variation of these two species, see Figure 6 in the manuscript. **Right panel: The relative average body proportions by species and by sex**.

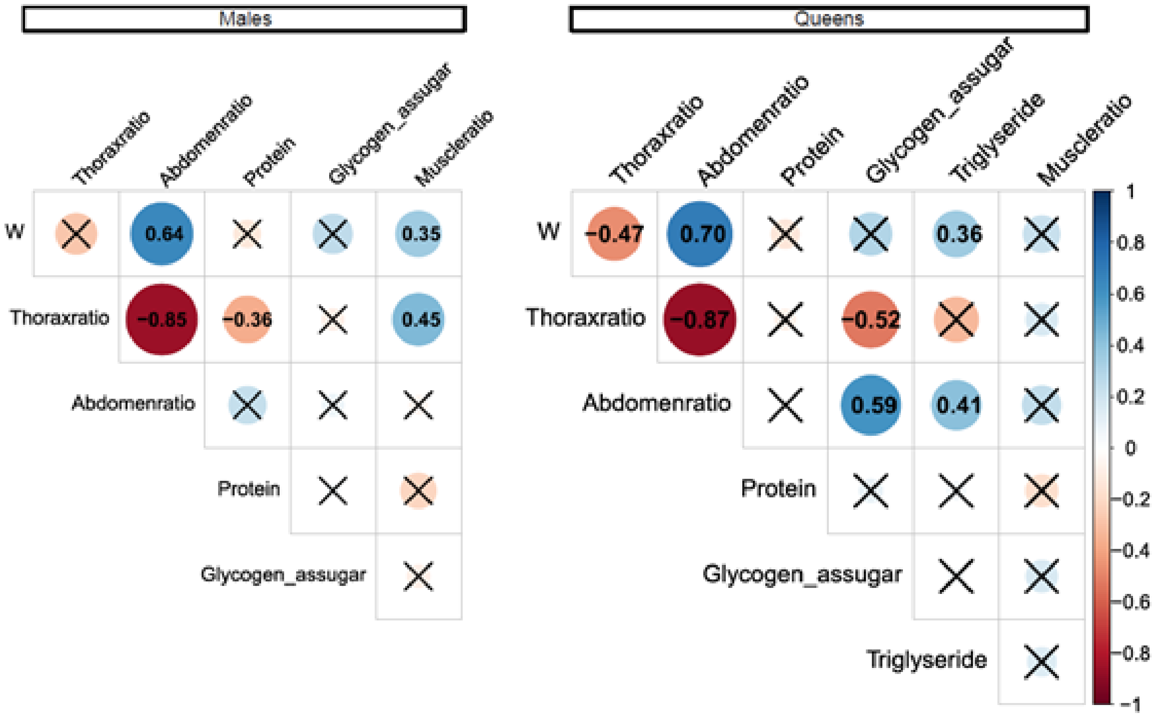


**Figure 2: Correlation coefficients among the morphological and biochemical variables**. Blue signifies positive correlations, red signifies negative correlations. X marks statistically non-significant correlations (p>0.01).

**Table 1: Differences in variances in body proportion and biochemical data with F test.** Social: comparison between monodomous and polydomous social structure, F values smaller than 1 indicate that the polydomous have a higher variance. Sex: comparison between males and queens, F value smaller than 1 indicates the queens have a higher variance.

| Measurement | Variance of data | |
|---|---|---|
| | Social | Sex |
| Total body weight | F = 1.89, p < 0.001 | F = 0.50, p < 0.001 |
| Abdomen ratio | F = 1.29, p = 0.001 | F = 0.37, p < 0.001 |
| Thorax ratio | F = 1.35, p < 0.001 | F = 0.38, p < 0.001 |
| Muscle ratio | F = 0.78, p = 0.04 | F = 1.16, p = 0.22 |
| Protein concentration | F = 0.66, p < 0.001 | F = 1.41, p = 0.003 |
| Glycogen concentration | F = 1.14, p = 0.25 | F = 0.45, p < 0.001 |
| Triglyceride concentration | F = 2.79, p < 0.001 | NA |

**Table 2 (next page): Analyses of the body proportions and biochemical measurements within *F. exsecta* and *F. pressilabris*.** The social organization (monodomous/polydomous), sex, and their interaction were used as fixed effects. In the cases when there is data only for a single sex (all biochemical measurements for *F. exsecta*, triglyceride for *F. pressilabris*), only the social organization is used as fixed effect. The nest was used as random effect. The Benjamini-Hochberg procedure (1995) was used to correct the parameter p-values to decrease the false discovery rate. Tests where the p-value is smaller than the BH critical value are considered significant. Non-significant results are presented in grey.

| | Response<br>Parameter names | | | LMM results | | |
|---|---|---|---|---|---|---|
| ***F. exsecta*** | **Total body mass** | **β** | **SE** | **t-value** | **p** | **BH** |
| | Intercept (Monodomous males) | 0.010 | 0.000 | 26.990 | <2E-16 | |
| | Polydomous | -0.005 | 0.001 | -8.041 | 0.000 | <0.001 |
| | Queens | 0.002 | 0.000 | 6.318 | 0.000 | <0.001 |
| | Interaction | 0.001 | 0.001 | 1.058 | 0.291 | 0.407 |
| | **Abdomen ratio** | | | | | |
| | Intercept (Monodomous males) | 0.378 | 0.005 | 82.091 | <2E-16 | |
| | Polydomous | 0.020 | 0.007 | 2.692 | 0.012 | 0.020 |
| | Queens | -0.033 | 0.005 | -6.387 | 0.000 | <0.001 |
| | Interaction | -0.001 | 0.013 | -0.103 | 0.918 | 0.972 |
| | **Thorax ratio** | | | | | |
| | Intercept (Monodomous males) | 0.537 | 0.004 | 130.694 | <2E-16 | |
| | Polydomous | -0.037 | 0.007 | -5.548 | 0.000 | <0.001 |
| | Queens | -0.047 | 0.004 | -10.536 | <2E-16 | <0.001 |
| | Interaction | 0.009 | 0.011 | 0.804 | 0.422 | 0.537 |
| | **Muscle ratio** | | | | | |
| | Intercept (Monodomous males) | 0.293 | 0.006 | 51.879 | <2E-16 | |
| | Polydomous | -0.051 | 0.010 | -4.862 | 0.000 | <0.001 |
| | Queens | -0.084 | 0.007 | -11.411 | <2E-16 | <0.001 |
| | Interaction | 0.043 | 0.018 | 2.348 | 0.021 | 0.033 |
| | **Protein concentration (males)** | | | | | |
| | Intercept (Monodomous) | 3.401 | 0.516 | 6.596 | 0.000 | |
| | Polydomous | -0.027 | 0.754 | -0.036 | 0.972 | 0.972 |
| | **Glycogen concentration (males)** | | | | | |
| | Intercept (Monodomous) | 0.886 | 0.157 | 5.637 | 0.000 | |
| | Polydomous | -0.150 | 0.230 | -0.654 | 0.521 | 0.608 |
| ***F. pressilabris*** | **Total body mass** | | | | | |
| | Intercept (Monodomous males) | 0.005 | 0.000 | 23.010 | <2E-16 | |
| | Polydomous | -0.001 | 0.000 | -2.487 | 0.020 | 0.093 |
| | Queens | -0.001 | 0.000 | -3.128 | 0.003 | 0.017 |
| | Interaction | 0.001 | 0.000 | 1.664 | 0.100 | 0.271 |
| | **Abdomen ratio** | | | | | |
| | Intercept (Monodomous males) | 0.374 | 0.007 | 52.347 | <2E-16 | |
| | Polydomous | 0.002 | 0.009 | 0.260 | 0.797 | 0.841 |
| | Queens | 0.003 | 0.012 | 0.265 | 0.793 | 0.841 |
| | Interaction | -0.032 | 0.015 | -2.195 | 0.033 | 0.105 |
| | **Thorax ratio** | | | | | |
| | Intercept (Monodomous males) | 0.515 | 0.006 | 82.562 | <2E-16 | |
| | Polydomous | -0.007 | 0.008 | -0.893 | 0.379 | 0.719 |
| | Queens | -0.095 | 0.011 | -8.898 | 0.000 | <0.001 |
| | Interaction | 0.029 | 0.013 | 2.273 | 0.028 | 0.105 |
| | **Muscle ratio** | | | | | |
| | Intercept (Monodomous males) | 0.219 | 0.012 | 19.080 | 0.000 | |
| | Polydomous | 0.016 | 0.016 | 0.995 | 0.329 | 0.695 |
| | Queens | -0.101 | 0.021 | -4.906 | 0.000 | <0.001 |
| | Interaction | 0.015 | 0.026 | 0.595 | 0.557 | 0.739 |
| | **Protein concentration** | | | | | |
| | Intercept (Monodomous males) | 3.445 | 0.319 | 10.802 | 0.000 | |
| | Polydomous | 0.228 | 0.452 | 0.505 | 0.617 | 0.739 |
| | Queens | -0.411 | 0.553 | -0.743 | 0.460 | 0.739 |
| | Interaction | -0.927 | 0.688 | -1.349 | 0.182 | 0.432 |
| | **Glycogen concentration** | | | | | |
| | Intercept (Monodomous males) | 0.482 | 0.165 | 2.919 | 0.006 | |
| | Polydomous | 0.129 | 0.228 | 0.566 | 0.576 | 0.739 |
| | Queens | -0.006 | 0.174 | -0.032 | 0.975 | 0.975 |
| | Interaction | 0.103 | 0.208 | 0.494 | 0.622 | 0.739 |
| | **Triglyceride concentration (queens)** | | | | | |
| | Intercept (Monodomous) | 0.305 | 0.159 | 1.920 | 0.084 | |
| | Polydomous | -0.132 | 0.195 | -0.678 | 0.513 | 0.739 |
