## Appendix 1 for "DISPERSAL BEHAVIOR RATHER THAN DISPERSAL MORPHOLOGY CREATES SOCIAL POLYMORPHISM IN FORMICA ANTS"

### APPENDIX 1: RESULTS FOR ALL MEASURED VARIABLES

Table 1 (next page): Results and sample sizes for all of the measured variables for each species and both sexes. For *F. exsecta* and *F. pressilabris*, the numbers presented here are calculated from their main populations. For measurements from the additional populations with differing social organization, see Table 2. The biochemical resources are presented as mass-corrected concentrations in TBE buffer (10 ul/ 1 mg) to allow comparisons among the differently sized species. The total mitochondria and myofibril areas are measured from TEM images with a total area of 69.2 $\mu m^2$. The male triglyceride measurements are shown in grey, as they are based on a very few individuals while most males had zero values.

| | | *F. pratensis* | | *F. aquilonia* | | *F. exsecta* | | *F. pressilabris* | | *F. fusca* | | *F. cinerea* | |
|---|---|---|---|---|---|---|---|---|---|---|---|---|---|
| | | **Monodomous** | | **Polydomous** | | **Monodomous** | | **Polydomous** | | **Monodomous** | | **Polydomous** | |
| | | ♂ | ♀ | ♂ | ♀ | ♂ | ♀ | ♂ | ♀ | ♂ | ♀ | ♂ | ♀ |
| | N ind | 148 | 153 | 135 | 142 | 125 | 112 | 87 | 71 | 69 | 93 | 127 | 74 |
| | N nest | 16 | 17 | 11 | 13 | 13 | 10 | 12 | 8 | 13 | 15 | 29 | 28 |
| **Weight (g)** | Mean | 0.0285 | 0.0388 | 0.0184 | 0.0255 | 0.0104 | 0.0124 | 0.0039 | 0.0040 | 0.0144 | 0.0225 | 0.0164 | 0.0232 |
| | SD | 0.0035 | 0.0046 | 0.0022 | 0.0019 | 0.0023 | 0.0013 | 0.0008 | 0.0004 | 0.0017 | 0.0044 | 0.0018 | 0.0026 |
| **Abdomen ratio** | Mean | 0.435 | 0.459 | 0.440 | 0.473 | 0.374 | 0.347 | 0.373 | 0.350 | 0.378 | 0.485 | 0.420 | 0.444 |
| | SD | 0.018 | 0.022 | 0.041 | 0.024 | 0.024 | 0.032 | 0.025 | 0.038 | 0.028 | 0.060 | 0.024 | 0.037 |
| **Thorax ratio** | Mean | 0.502 | 0.427 | 0.496 | 0.415 | 0.540 | 0.489 | 0.509 | 0.443 | 0.548 | 0.411 | 0.514 | 0.447 |
| | SD | 0.016 | 0.019 | 0.038 | 0.022 | 0.023 | 0.024 | 0.025 | 0.028 | 0.028 | 0.048 | 0.023 | 0.031 |
| | N ind | 53 | 62 | 58 | 50 | 54 | 47 | 35 | 29 | 30 | 42 | 60 | 32 |
| | N nest | 11 | 14 | 11 | 10 | 13 | 10 | 10 | 7 | 13 | 15 | 26 | 19 |
| **Muscle ratio** | Mean | 0.29 | 0.20 | 0.27 | 0.20 | 0.29 | 0.21 | 0.23 | 0.15 | 0.31 | 0.22 | 0.30 | 0.25 |
| | SD | 0.01 | 0.01 | 0.03 | 0.02 | 0.03 | 0.03 | 0.04 | 0.04 | 0.03 | 0.03 | 0.03 | 0.02 |
| | N ind | 72 | 69 | 51 | 71 | 55 | 41 | 40 | 34 | 26 | 40 | 58 | 31 |
| | N nest | 11 | 14 | 15 | 14 | 11 | 7 | 11 | 8 | 10 | 12 | 21 | 15 |
| **Protein (mg/ml)** | Mean | 3.252 | 1.729 | 4.299 | 3.124 | 3.504 | 2.607 | 3.673 | 2.319 | 2.573 | 1.810 | 3.827 | 1.830 |
| | SD | 1.804 | 1.099 | 2.448 | 2.401 | 1.773 | 1.474 | 1.831 | 0.585 | 1.865 | 1.401 | 1.851 | 1.134 |
| **Glycogen (mg/ml)** | Mean | 1.663 | 1.314 | 1.078 | 2.155 | 0.889 | 1.231 | 0.913 | 0.343 | 0.526 | 2.413 | 0.506 | 1.417 |
| | SD | 1.014 | 0.811 | 0.864 | 1.178 | 0.721 | 1.272 | 0.726 | 0.260 | 0.566 | 1.824 | 0.574 | 1.350 |
| **Free glucose (mg/ml)** | Mean | 1.390 | 0.720 | 1.410 | 0.971 | 1.167 | 0.680 | 1.029 | 0.231 | 0.507 | 0.353 | 0.370 | 1.008 |
| | SD | 0.559 | 0.443 | 0.632 | 0.494 | 0.753 | 0.448 | 0.612 | 0.176 | 0.546 | 0.258 | 0.462 | 0.709 |
| **Triglyceride (mg/ml)** | Mean | 0.002 | 0.558 | 0.003 | 0.288 | 0.012 | 0.093 | 0.035 | 0.213 | 0.000 | 0.550 | 0.037 | 0.452 |
| | SD | 0.013 | 0.520 | 0.011 | 0.292 | 0.069 | 0.228 | 0.077 | 0.251 | 0.000 | 0.422 | 0.168 | 0.372 |
| | N ind | 9 | 9 | 9 | 9 | 9 | 9 | 9 | 9 | 9 | 9 | 9 | 9 |
| | N nest | 3 | 3 | 3 | 3 | 3 | 3 | 3 | 3 | 3 | 3 | 3 | 3 |
| **Total mitochondria area (um2)** | Mean | 26.906 | 27.426 | 27.974 | 29.980 | 26.167 | 28.037 | 27.704 | 28.040 | 30.168 | 27.545 | 25.487 | 27.253 |
| | SD | 3.989 | 3.123 | 3.609 | 2.547 | 3.354 | 4.486 | 3.706 | 3.058 | 3.793 | 3.786 | 3.302 | 4.280 |
| **Mitochondrium profile area (um2)** | Mean | 0.887 | 0.754 | 0.694 | 0.992 | 0.957 | 0.997 | 1.028 | 0.925 | 1.319 | 0.876 | 0.736 | 0.715 |
| | SD | 0.532 | 0.480 | 0.463 | 0.584 | 0.686 | 0.649 | 0.635 | 0.585 | 0.844 | 0.608 | 0.470 | 0.505 |
| **Mitochondrium profile degraded area (um2)** | Mean | 0.020 | 0.014 | 0.033 | 0.023 | 0.031 | 0.046 | 0.068 | 0.042 | 0.063 | 0.022 | 0.022 | 0.036 |
| | SD | 0.078 | 0.051 | 0.095 | 0.067 | 0.097 | 0.152 | 0.205 | 0.157 | 0.188 | 0.095 | 0.089 | 0.144 |
| **Mitochondrium profile shape** | Mean | 0.272 | 0.305 | 0.283 | 0.298 | 0.311 | 0.320 | 0.294 | 0.302 | 0.337 | 0.337 | 0.281 | 0.273 |
| | SD | 0.145 | 0.154 | 0.149 | 0.149 | 0.157 | 0.159 | 0.158 | 0.153 | 0.164 | 0.157 | 0.144 | 0.142 |
| **Total myofibril area (um2)** | Mean | 30.322 | 33.325 | 32.645 | 31.118 | 28.771 | 30.174 | 28.424 | 31.607 | 26.770 | 30.922 | 33.321 | 30.180 |
| | SD | 3.208 | 3.268 | 3.108 | 2.564 | 4.101 | 4.737 | 3.558 | 4.347 | 3.047 | 3.950 | 3.160 | 4.665 |
| **Myofibril profile diameter (um)** | Mean | 1.660 | 1.741 | 1.619 | 1.505 | 1.649 | 1.549 | 1.459 | 1.559 | 1.444 | 1.529 | 1.631 | 1.571 |
| | SD | 0.302 | 0.316 | 0.256 | 0.231 | 0.279 | 0.324 | 0.250 | 0.257 | 0.289 | 0.269 | 0.268 | 0.241 |
| **CV of distances among myofibrils** | Mean | 0.237 | 0.218 | 0.236 | 0.241 | 0.228 | 0.233 | 0.257 | 0.242 | 0.252 | 0.232 | 0.228 | 0.230 |
| | SD | 0.094 | 0.087 | 0.077 | 0.069 | 0.080 | 0.086 | 0.076 | 0.085 | 0.059 | 0.077 | 0.071 | 0.071 |

Table 2: Results and sample sizes for both of the social organizations of *F. exsecta* and *F. pressilabris*. The biochemical resources are presented as mass-corrected concentrations in TBE buffer (10 ul/ 1 mg) to allow comparisons among the differently sized species. The male triglyceride measurements are shown in grey, as they are based on a very few individuals while most males had zero values.

| | | ***F. exsecta*** | | | | ***F. pressilabris*** | | | |
|---|---|---|---|---|---|---|---|---|---|
| | | **Monodomous** | | **Polydomous** | | **Monodomous** | | **Polydomous** | |
| | | ♂ | ♀ | ♂ | ♀ | ♂ | ♀ | ♂ | ♀ |
| | N ind | 125 | 112 | 65 | 7 | 78 | 29 | 87 | 71 |
| | N nest | 13 | 10 | 10 | 1 | 9 | 4 | 12 | 8 |
| **Weight (g)** | Mean | 0.0104 | 0.0124 | 0.0054 | 0.0099 | 0.0045 | 0.0043 | 0.0039 | 0.0040 |
| | SD | 0.0023 | 0.0013 | 0.0014 | 0.0006 | 0.0008 | 0.0004 | 0.0008 | 0.0004 |
| **Abdomen ratio** | Mean | 0.374 | 0.347 | 0.397 | 0.370 | 0.373 | 0.374 | 0.373 | 0.350 |
| | SD | 0.024 | 0.032 | 0.029 | 0.032 | 0.029 | 0.037 | 0.025 | 0.038 |
| **Thorax ratio** | Mean | 0.540 | 0.489 | 0.500 | 0.457 | 0.517 | 0.422 | 0.509 | 0.443 |
| | SD | 0.023 | 0.024 | 0.027 | 0.025 | 0.030 | 0.027 | 0.025 | 0.028 |
| | N ind | 54 | 47 | 19 | 7 | 35 | 14 | 35 | 29 |
| | N nest | 13 | 10 | 9 | 1 | 8 | 4 | 10 | 7 |
| **Muscle ratio** | Mean | 0.293 | 0.210 | 0.243 | 0.200 | 0.217 | 0.116 | 0.235 | 0.151 |
| | SD | 0.032 | 0.025 | 0.046 | 0.035 | 0.061 | 0.031 | 0.038 | 0.036 |
| | N ind | 55 | 41 | 47 | 0 | 50 | 15 | 40 | 34 |
| | N nest | 11 | 7 | 10 | 0 | 12 | 4 | 12 | 8 |
| **Protein (mg/ml)** | Mean | 3.504 | 2.607 | 3.086 | NA | 3.504 | 2.848 | 3.673 | 2.319 |
| | SD | 1.773 | 1.474 | 2.169 | NA | 0.950 | 1.262 | 1.831 | 0.585 |
| **Glycogen (mg/ml)** | Mean | 0.889 | 1.231 | 0.835 | NA | 0.523 | 0.429 | 0.913 | 0.343 |
| | SD | 0.721 | 1.272 | 0.489 | NA | 0.634 | 0.311 | 0.726 | 0.260 |
| **Free glucose (mg/ml)** | Mean | 1.164 | 0.676 | 1.195 | NA | 1.135 | 0.616 | 1.029 | 0.223 |
| | SD | 0.757 | 0.456 | 0.562 | NA | 0.582 | 0.208 | 0.612 | 0.187 |
| **Triglyceride (mg/ml)** | Mean | 0.012 | 0.093 | 0.015 | NA | 0.047 | 0.272 | 0.035 | 0.213 |
| | SD | 0.069 | 0.228 | 0.067 | NA | 0.088 | 0.376 | 0.077 | 0.251 |
