## Appendix 3 for "DISPERSAL BEHAVIOR RATHER THAN DISPERSAL MORPHOLOGY CREATES SOCIAL POLYMORPHISM IN FORMICA ANTS"

### APPENDIX 3: MICROSCOPICAL MUSCLE STRUCTURES

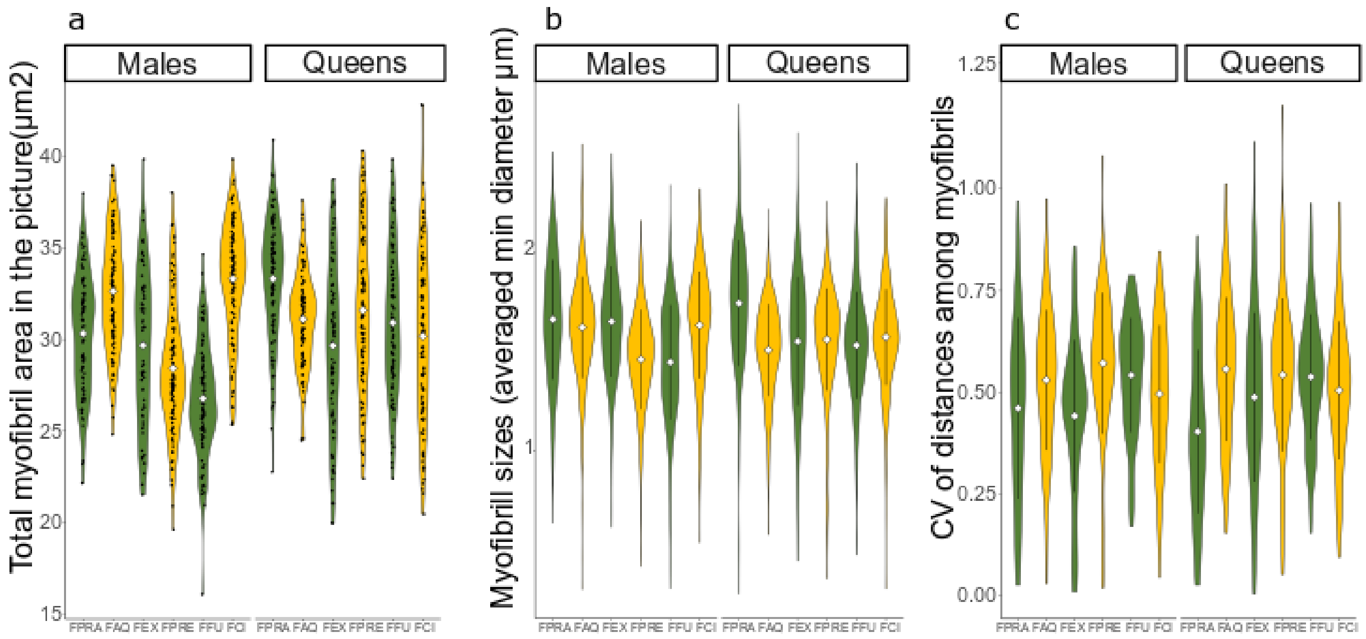


**Figure 1. Results of the myofibril measurements**. a) Total myofibril area per TEM image (image area=69.2 µm$^2$) with all data points, b) size of individual myofibrils as diameters with mean and standard deviation, c) organization of myofibrils per TEM image as coefficient of variation, with mean and standard deviation. All data are visualized as density plots (violin plots), the mean is visualized with a diamond shape. The species are abbreviated as follows: FPRA = *F. pratensis*, FAQ = *F. aquilonia*, FEX = *F.exsecta*, FPRE = *F. pressilabris*, FFU = *F. fusca*, FCI = *F.cinerea*. The social organization of each species is highlighted with color: dark green = monodomous, yellow = polydomous.

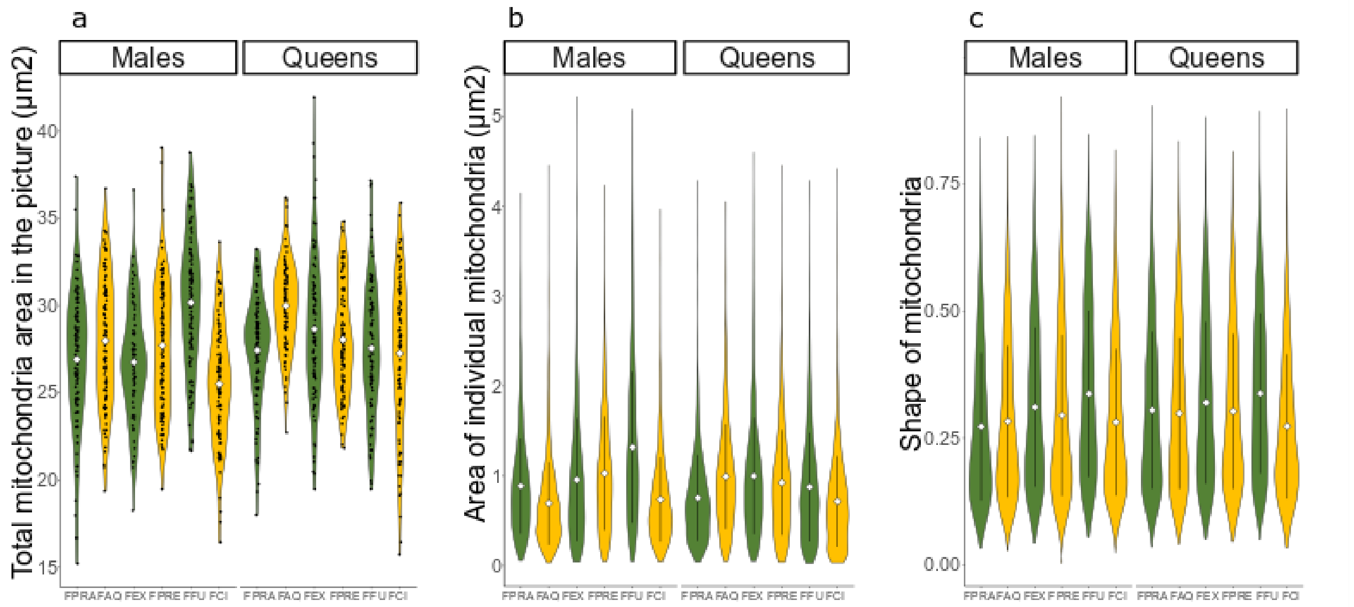


**Figure 2. Results of the mitochondria measurements. a**) Total mitochondria area per TEM image (image area=69.2 µm$^2$) with all data points, b) area of individual mitochondria with mean and standard deviation, c) mitochondrion profile shape, with mean and standard deviation. 0= round, longer shapes, especially > 0.5 indicate fused and tubular, strong mitochondria. All data are visualized as density plots, the mean is visualized with a diamond shape. The species are abbreviated as follows: FPRA = *F. pratensis*, FAQ = *F. aquilonia*, FEX = *F.exsecta*, FPRE = *F. pressilabris*, FFU = *F. fusca*, FCI = *F.cinerea*. The social organization of each species is highlighted with color: dark green = monodomous, yellow = polydomous.

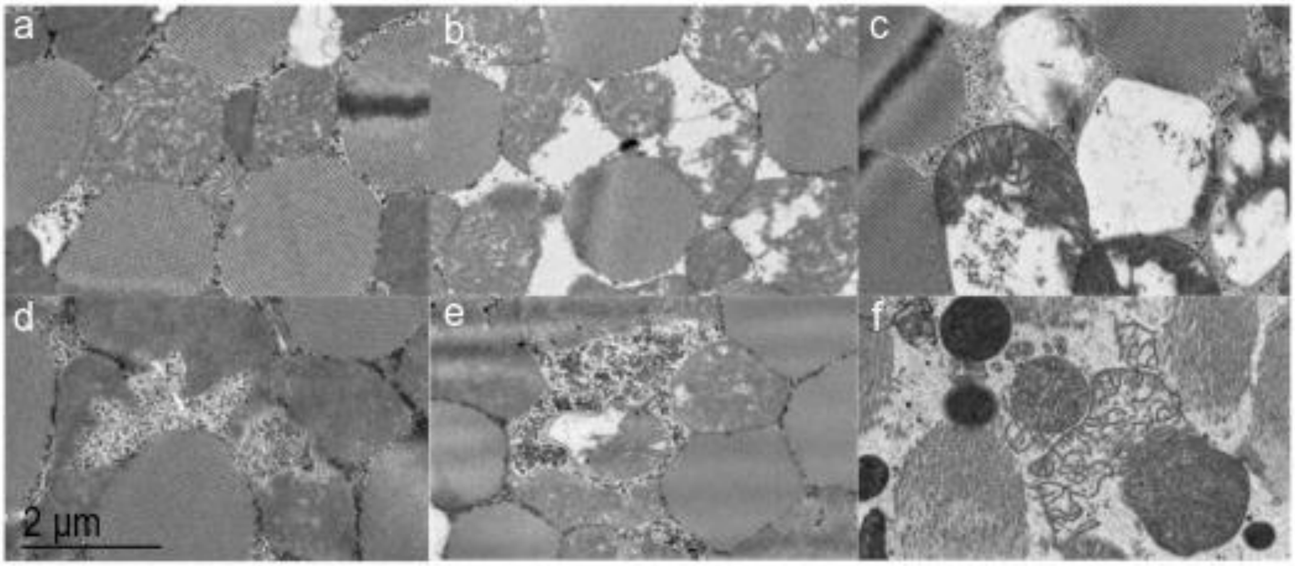


**Figure 3. Examples of mitochondria structures that get values for degradation.** a) Slightly loose cristae structure unlike the usually denser cristae in our specimens – this is not true degradation but gives low values with our thresholding, b&c) empty area within mitochondria membranes, d&e) empty area filled with glycogen inside mitochondria membranes, f) extremely degraded mitochondria with broken membranes. Small values of degradation are common in our specimens and likely normal with no effects for flight ability. High values (>10% of the mitochondria profile area) are likely related to decrease of muscle function.

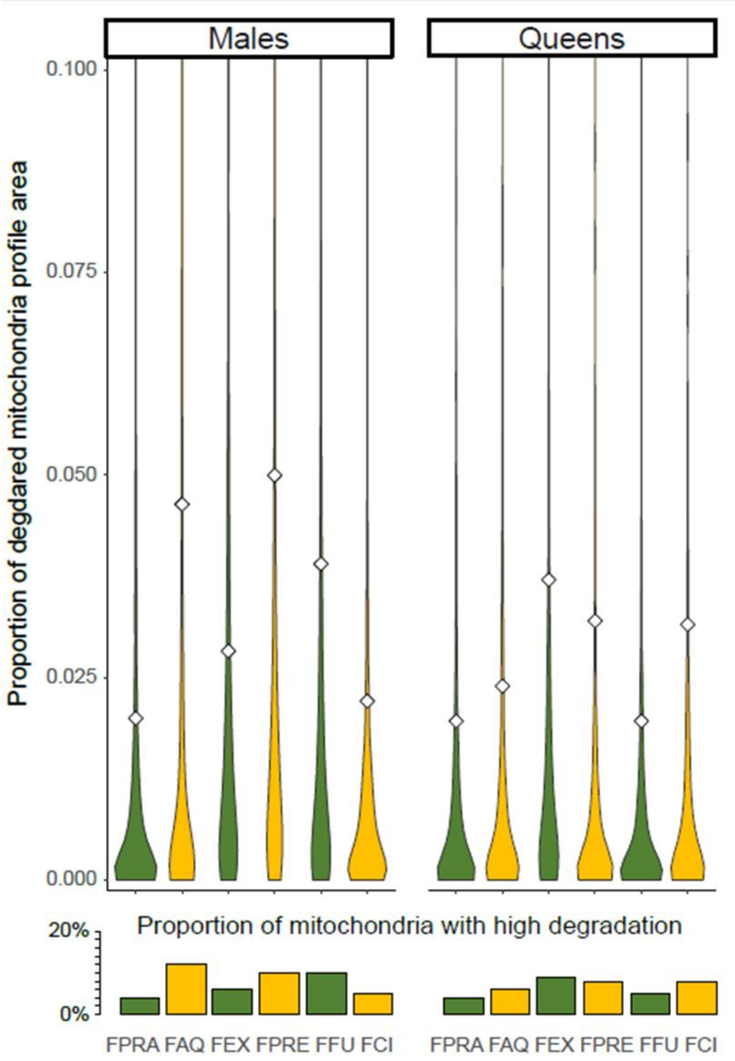


**Figure 4. Mitochondria degradation for each species and both sexes.** Upper panel shows the proportion of degraded area per total area of mitochondria profiles. Most mitochondria have low values for degradation (mean= 0.03). The y-axis of the upper panel is cut at 0.1, and the lower panel shows the proportion of data left out from the upper panel: the percentage of mitochondria having higher than 0.1 proportion of degraded area. Upper panel data are visualized as kernel density plots, the mean is visualized with a diamond shape. The species are abbreviated as follows: FPRA = *F. pratensis*, FAQ = *F. aquilonia*, FEX = *F.exsecta*, FPRE = *F. pressilabris*, FFU = *F. fusca*, FCI = *F.cinerea*. The social organization of each species is highlighted with color: dark green = monodomous, yellow = polydomous.

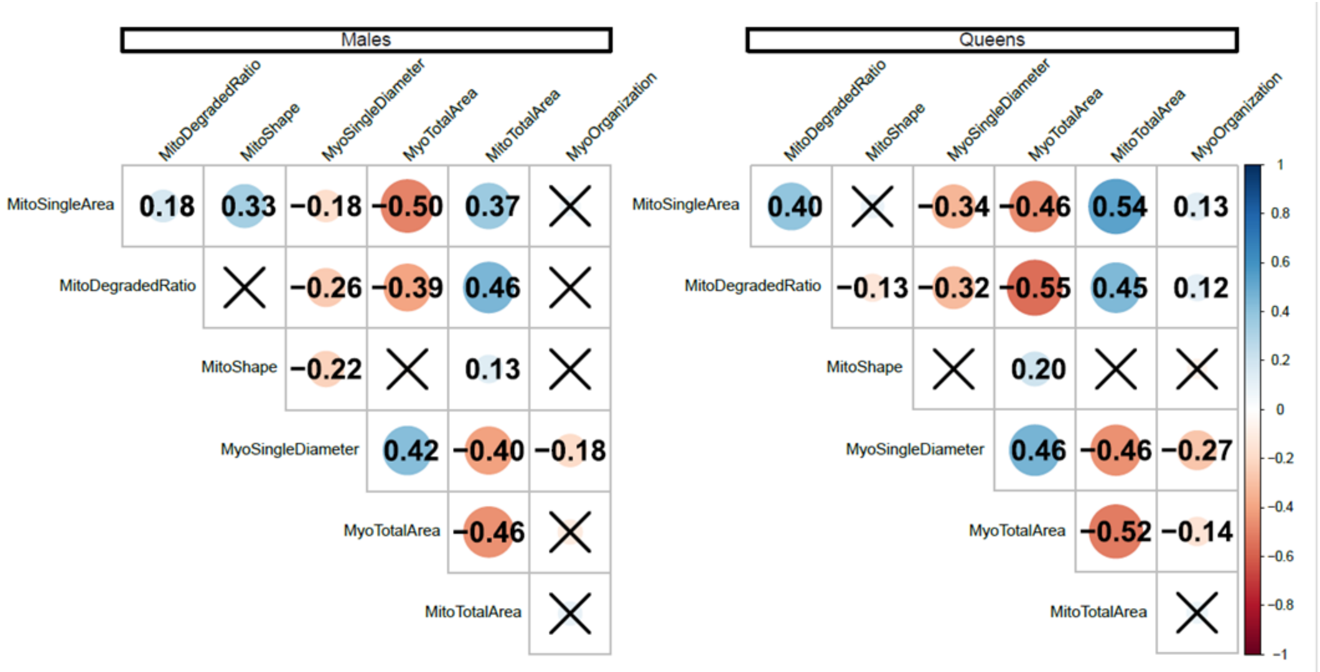


**Figure 5. Correlation coefficients among the microscopical muscle structure variables.** Blue signifies positive correlations, red signifies negative correlations. X marks statistically non-significant correlations (p>0.01).

**Table 1: Analyses of the microscopical muscle structure measurements**. In all of the analyses, the social organization (monodomous/polydomous), sex, and their interaction were used as fixed effects, and the species, nest and individual identity as nested random effects. For the analyses of organelle profiles also the image was used as a random effect, nested within individual. The Benjamini-Hochberg procedure (1995) was used to correct the parameter p-values to decrease the false discovery rate. Tests where the p-value is smaller than the BH critical value are considered significant. Non-significant results are presented in grey.

| | **Response**<br>Parameter names | **β** | **SE** | **z/t-value** | **p** | **BH** |
|---|---|---|---|---|---|---|
| **Mitochondria** | **Total area** | | | **LMM** | | |
| | Intercept (Monodomous males) | 27.72407 | 0.74202 | 37.363 | 1.05E-08 | |
| | Polydomous | -0.66302 | 1.04898 | -0.632 | 0.549 | 0.721 |
| | Queens | -0.08522 | 0.68776 | -0.124 | 0.902 | 0.947 |
| | Interaction | 1.44588 | 0.97661 | 1.481 | 0.15 | 0.394 |
| | **Area of profiles** | | | **GLMM (gamma)** | | |
| | Intercept (Monodomous males) | 1.00035 | 0.09196 | 10.878 | <2E-16 | |
| | Polydomous | 0.28438 | 0.12995 | 2.188 | 0.0286 | 0.200 |
| | Queens | 0.18921 | 0.1079 | 1.754 | 0.0795 | 0.278 |
| | Interaction | -0.27134 | 0.15286 | -1.775 | 0.0759 | 0.278 |
| | **Shape of profiles** | | | **GLMM (gamma)** | | |
| | Intercept (Monodomous males) | 3.4187 | 0.1089 | 31.391 | <2E-16 | |
| | Polydomous | 0.192 | 0.1536 | 1.25 | 0.211 | 0.443 |
| | Queens | -0.1756 | 0.1184 | -1.483 | 0.138 | 0.394 |
| | Interaction | 0.1182 | 0.1678 | 0.704 | 0.481 | 0.678 |
| | **Presense of high (>10) degradation** | | | **GLMM (binomial)** | | |
| | Intercept (Monodomous males) | -3.21257 | 0.22436 | -14.319 | <2E-16 | |
| | Polydomous | 0.22477 | 0.31303 | 0.718 | 0.473 | 0.678 |
| | Queens | -0.12972 | 0.31424 | -0.413 | 0.68 | 0.840 |
| | Interaction | 0.08255 | 0.44317 | 0.186 | 0.852 | 0.942 |
| **Myofibrils** | **Total area** | | | **LMM** | | |
| | Intercept (Monodomous males) | 28.5957 | 0.9097 | 31.436 | 9.80E-10 | |
| | Polydomous | 2.8304 | 1.2862 | 2.201 | 0.05865 | 0.278 |
| | Queens | 2.8813 | 1.0079 | 2.859 | 0.00795 | 0.167 |
| | Interaction | -3.371 | 1.4291 | -2.359 | 0.0255 | 0.200 |
| | **Diameter of profiles** | | | **LMM** | | |
| | Intercept (Monodomous males) | 1.62538 | 0.05604 | 29.003 | 1.70E-07 | |
| | Polydomous | -0.02046 | 0.07922 | -0.258 | 0.805 | 0.939 |
| | Queens | 0.03296 | 0.04645 | 0.71 | 0.484 | 0.678 |
| | Interaction | -0.06492 | 0.06586 | -0.986 | 0.333 | 0.636 |
| | **CV of distances among myofibrils** | | | **LMM** | | |
| | Intercept (Monodomous males) | 0.2395166 | 0.0069459 | 34.483 | 1.01E-12 | |
| | Polydomous | 0.0003942 | 0.0096782 | 0.041 | 0.968 | 0.968 |
| | Queens | -0.0118959 | 0.0087021 | -1.367 | 0.183 | 0.427 |
| | Interaction | 0.009038 | 0.01219 | 0.741 | 0.465 | 0.678 |
